# Reduction of a Stochastic Model of Gene Expression: Lagrangian Dynamics Gives Access to Basins of Attraction as Cell Types and Metastabilty

**DOI:** 10.1101/2020.09.04.283176

**Authors:** Elias Ventre, Thibault Espinasse, Charles-Edouard Bréhier, Vincent Calvez, Thomas Lepoutre, Olivier Gandrillon

## Abstract

Differentiation is the process whereby a cell acquires a specific phenotype, by differential gene expression as a function of time. This is thought to result from the dynamical functioning of an underlying Gene Regulatory Network (GRN). The precise path from the stochastic GRN behavior to the resulting cell state is still an open question. In this work we propose to reduce a stochastic model of gene expression, where a cell is represented by a vector in a continuous space of gene expression, to a discrete coarse-grained model on a limited number of cell types. We develop analytical results and numerical tools to perform this reduction for a specific model characterizing the evolution of a cell by a system of piecewise deterministic Markov processes (PDMP). Solving a spectral problem, we find the explicit variational form of the rate function associated to a large deviations principle, for any number of genes. The resulting Lagrangian dynamics allows us to define a deterministic limit of which the basins of attraction can be identified to cellular types. In this context the quasipotential, describing the transitions between these basins in the weak noise limit, can be defined as the unique solution of an Hamilton-Jacobi equation under a particular constraint. We develop a numerical method for approximating the coarse-grained model parameters, and show its accuracy for a symmetric toggle-switch network. We deduce from the reduced model an approximation of the stationary distribution of the PDMP system, which appears as a Beta mixture. Altogether those results establish a rigorous frame for connecting GRN behavior to the resulting cellular behavior, including the calculation of the probability of jumps between cell types.

## Introduction

Differentiation is the process whereby a cell acquires a specific phenotype, by differential gene expression as a function of time. Measuring how gene expression changes as differentiation proceeds is therefore of essence to understand this process. Advances in measurement technologies now allow to obtain gene expression levels at the single cell level. It offers a much more accurate view than population-based measurements, that has been obscured by mean population-based averaging [1], [2]. It has been established that there is a high cell-to-cell variability in gene expression, and that this variability has to be taken into account when investigating a differentiation process at the single-cell level [3], [4], [5], [6], [7], [8], [9], [10], [11].

A popular vision of the cellular evolution during differentiation, introduced by Waddington in [12], is to compare cells to marbles following probabilistic trajectories, as they roll through a developmental landscape of ridges and valleys. These trajectories are represented in the gene expression space: a cell can be described by a vector, each coordinate of which represents the expression of a gene [13], [14]. Thus, the state of a cell is characterized by its position in the gene expression space, *i.e* its specific level for all of its expressed genes. This landscape is often considered to be shaped by the underlying gene regulatory network (GRN), the behavior of which can be influenced by many factors, such as proliferation or cell-to-cell communication. Theoretically, the number of states a cell can take is equal to the number of possible combination of protein quantities associated to each gene. This number is potentially huge [15]. But metastability seems inherent to cell differentiation processes, as evidenced by limited number of existing cellular phenotypes [16], [17], providing a rationale for dimension reduction approaches [18]. Indeed, since [19] and [20], many authors have identified cell types with the basins of attraction of a dynamical system modeling the differentiation process, although the very concept of “cell type” has to be interrogated in the era of single-cell omics [21].

Adapting this identification for characterizing metastability in the case of stochastic models of gene expression has been studied mostly in the context of stochastic diffusion processes [22], [23], [24], but also for stochastic hybrid systems [25]. In the weak noise limit, a natural development of this analysis consists in describing the transitions between different macrostates within the large deviations framework [26], [27].

We are going to apply this strategy for a piecewise-deterministic Markov process (PDMP) describing GRN dynamics within a single cell, introduced in [28], which corresponds accurately to the non-Gaussian distribution of single–cell gene expression data. Using the work of [29], the novelty of this article is to provide analytical results for characterizing the metastable behavior of the model for any number of genes, and to combine them with a numerical analysis for performing the reduction of the model in a coarse-grained discrete process on cell types. We detail the model in Section 1, and we present in Section 2 how the reduction of this model in a continuous-time Markov chain on cell types allows to characterize the notion of metastability. For an arbitrary network, we provide in Section 3.1 a numerical method for approximating each transition rate of this coarse-grained model, depending on the probability of a rare event. In Section 3.2, we show that this probability is linked to a large deviations principle. The main contribution of this article is to derive in Section 4.1 the explicit variational form of the rate function associated to a Large deviations principle (LDP) for this model. We discuss in Sections 4.2 and 4.3 the conditions for which a unique quasipotential exists and allows to describe transitions between basins. We replace in Section 4.4 these results in the context of studying metastability. Finally, we apply in Section 5 the general results to a toggle-switch network. We also discuss in Section 6.1 some notions of energy associated to the LDP and we propose in Section 6.2 a non-Gaussian mixture model for approximating proteins distribution.

## 1 Model description

The model which is used throughout this article is based on a hybrid version of the well-established two-state model of gene expression [30], [31]. A gene is described by the state of the promoter, which can be {*on, off*}. If the state of the promoter is *on*, mRNAs are transcripted and translated into proteins, which are considered to be produced at a rate *s*. If the state of the promoter is *off*, only degradation of proteins occurs at a rate *d* (see Figure 1). *k*_*on*_ and *k*_*off*_ denote the exponential rates of transition between the states *on* and *off*. This model is a reduction of a mechanistic model including both mRNA and proteins, which is described in Appendix A.

**Figure 1:**
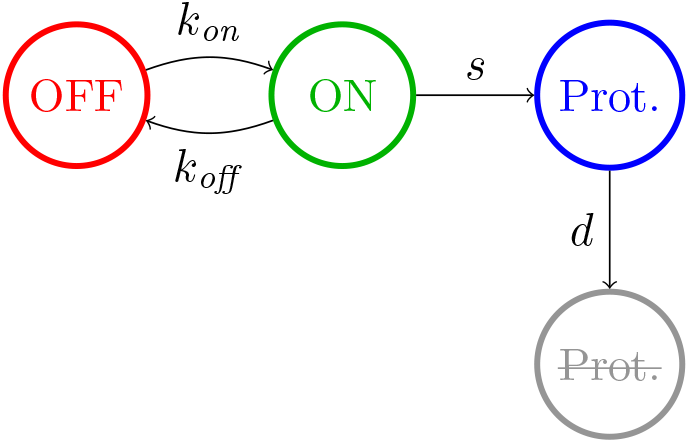
Simplified two-states model of gene expression [28], [31].

Neglecting the molecular noise associated to proteins quantity, we obtain the hybrid model:

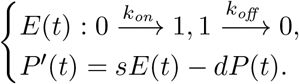

where *E*(*t*) denotes the promoter state, *P* (*t*) denotes the protein concentration at time *t*, and we identify the state *off* with 0, the state *on* with 1.

The key idea for studying a GRN is to embed this two-states model into a network. Denoting the number of genes by *n*, the vector (*E, P*) describing the process is then of dimension 2*n*. The jump rates for each gene *i* are expressed in terms of two specific functions *k*_*on,i*_ and *k*_*off,i*_. To take into account the interactions between the genes, we consider that for all *i* = 1, · · ·, *n*, *k*_*on,i*_ is a function which depends on the full vector *P* via the GRN, represented by a matrix Θ of size *n*. We assume that *k*_*on,i*_ is upper and lower bounded by a positive constant for all *i*. The function is chosen such that if gene *i* activates gene *j*, then 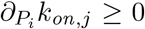. For the sake of simplicity, we consider that *k*_*off,i*_ does not depend on the protein level.

We introduce a typical time scale 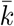 for the rates of promoters activation *k*_*on,i*_, and a typical time scale 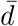 for the rates of proteins degradation. Then, we define the scaling factor 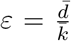 which characterizes the difference in dynamics between two processes: 1. gene bursting dynamics and 2. protein dynamics. It is generally considered that promoter switches are fast with respect to protein dynamics, *i.e* that *ε* ≪ 1, at least for eukaryotes [32]. Driven by biological considerations, we will consider values of *ε* smaller than 1/5 (see Appendix A).

We then rescale the time of the process by 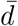. We also rescale the quantities *k*_*on,i*_ and *k*_*off,i*_ by 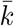, and *d*_*i*_ by 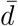, for any gene *i*, in order to simplify the notations. Finally, the parameters *s*_*i*_ can be removed by a simple rescaling of the protein concentration *P*_*i*_ for every gene by its equilibrium value when *E*_*i*_ = 1 (see [28] for more details). We obtain a reduced dimensionless PDMP system modeling the expression of *n* genes in a single cell:

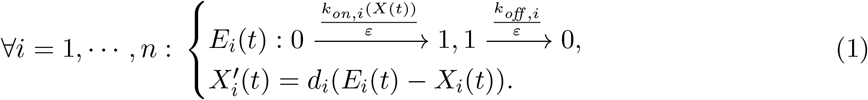

Here, *X* describes the protein vector in the renormalized gene expression space Ω := (0, 1)^*n*^ and *E* describes the promoters state, in *P*_*E*_ := {0, 1}^*n*^. We will refer to this model, that we will use throughout this article, as the PDMP system.

As card(*P*_*E*_) = 2^*n*^, we can write the joint probability density *u*(*t, e, x*) of (*E*_*t*_, *X*_*t*_) as a 2^*n*^-dimensional vector 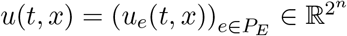. The master equation on *u* can be written:

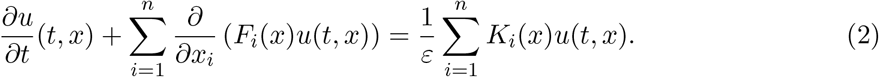

For all *i* = 1, · · ·, *n*, for all *x* ∈ Ω, *F*_*i*_(*x*) and *K*_*i*_(*x*) are matrices of size 2^*n*^. Each *F*_*i*_ is diagonal, and the term on a line associated to a promoter state *e* corresponds to the drift of gene *i*: *d*_*i*_(*e*_*i*_ − *x*_*i*_). *K*_*i*_ is not diagonal: each state *e* is coupled with every state *e*’ such that only the coordinate *e*_*i*_ changes in *e*, from 1 to 0 or conversely. Each of these matrices can be expressed as a tensorial product of (*n* − 1) two-dimensional identity matrices with a two-dimensional matrix corresponding to the operator associated to an isolated gene:

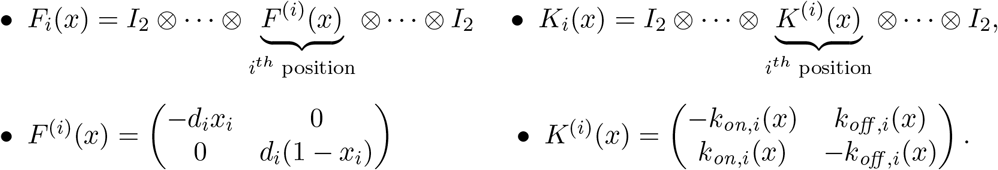

We detail in Appendix B the case of *n* = 2 for a better understanding of this tensorial expression.

## 2 Model reduction in the small noise limit

### 2.1 Deterministic approximation

The model (1) describes the promoter state of every gene *i* at every time as a Bernoulli random variable. We use the biological fact that promoter switches are frequent compared to protein dynamic, *i.e ε* < 1 with the previous notations. When *ε* ≪ 1, we can approximate the conditional distribution of the promoters knowing proteins, *ρ*, by its quasistationary approximation 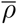:

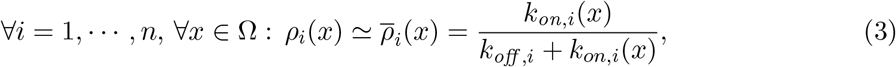

which is derived from the stationary distribution of the Markov chain on the promoters states, defined for a given value of the protein vector *X* = *x* by the matrix 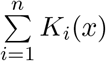 (see [33], [34]). Thus, the PDMP model (1) can be coarsely approximated by a system of ordinary differential equations:

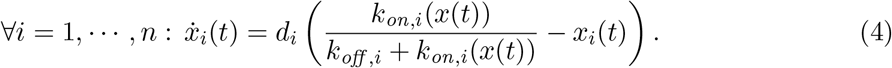

Intuitively, these trajectories correspond to the mean behaviour of a cell in the weak noise limit, *i.e* when promoters jump much faster than proteins concentration changes. More precisely, a random path 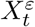 converges in probability to a trajectory *ϕ*_*t*_ solution of the system (4), when *ε* → 0 [35]. The diffusion limit, which keeps a residual noise scaled by 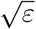, can also be rigorously derived from the PDMP system [36], which is detailed in Appendix C.1.

In the sequel, we assume that every limit set of a trajectory solution of the system (4) as *t* → +∞ is reduced to a single equilibrium point, described by one of the solutions of:

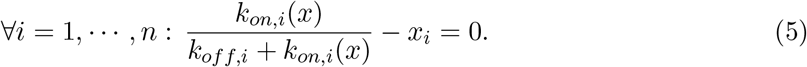

Note that the condition above strongly depends on the interaction functions {*k*_*on,i*_}_*i*=1,···,*n*_. Alternatively speaking, in this work we rule out the existence of attractive limit cycles or more complicated orbits. We also assume that the closure of the basins of attraction which are associated to the stable equilibria of the system (5) covers the gene expression space Ω.

Without noise, the fate of a cell trajectory is fully characterized by its initial state *x*_0_. Generically, it converges to the attractor of the basin of attraction it belongs to, which is a single point by assumption. However, noise can modify the deterministic trajectories in at least two ways. First, in short times, a stochastic trajectory can deviate significantly from the deterministic one. In the case of a single, global, attractor, the deterministic system generally allows to retrieve the global dynamics of the process, *i.e* the equilibrium and the order of convergence between the different genes, for realistic *ε* (see Figure 2).

**Figure 2:**
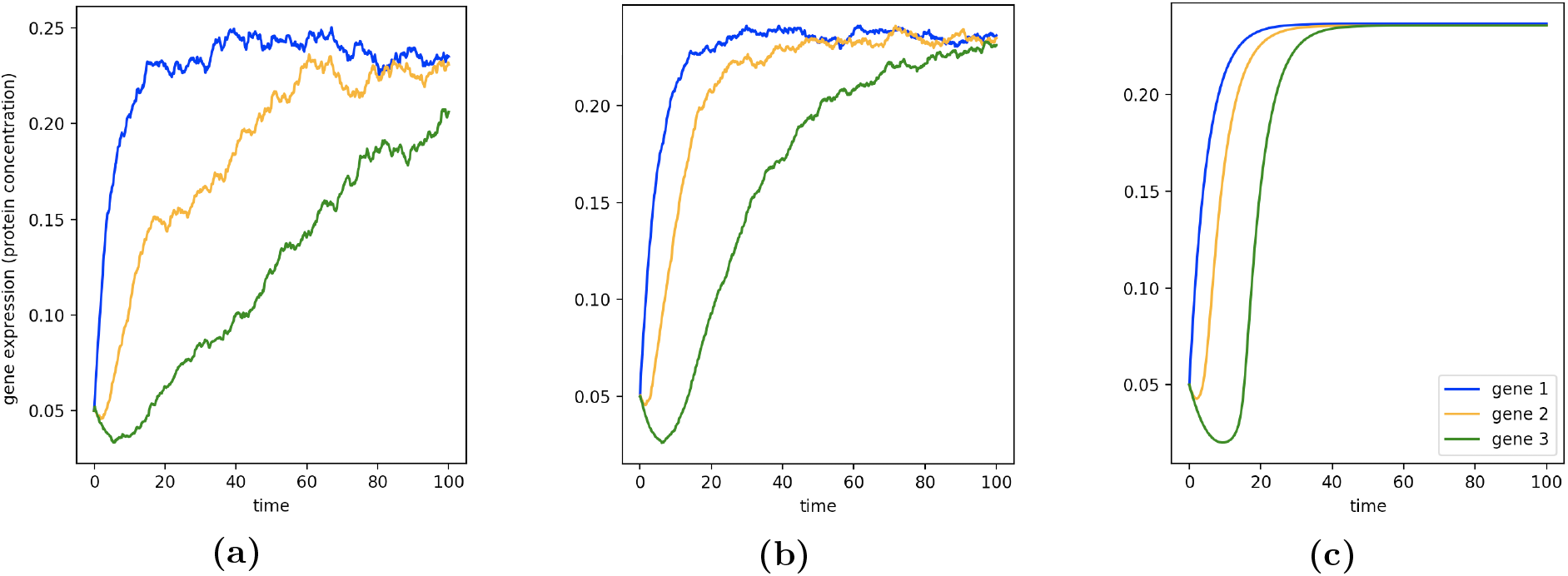
Comparison between the average on 100 simulated trajectories with *ε* = 1/7 (2a), *ε* = 1/30 (2b) and the trajectories generated by the deterministic system (2c) for a single pathway network: gene 1 → gene 2 → gene 3.

Second, in long times, stochastic dynamics can even push the trajectory out of the basin of attraction of one equilibrium state to another one, changing radically the fate of the cell. These transitions cannot be catched by the deterministic limit, and happen on a time scale which is expected to be of the order of 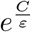 (owing to a Large deviations principle studied below), where *C* is an unknown constant depending on the basins. In Figure 3a, we illustrate this situation for a toggle-switch network of two genes. We observe possible transitions between three basins of attraction. Two examples of random paths, the stochastic evolution of promoters and proteins along time, are represented in Figures 3b and 3c for different values of *ε*. All the details on the interaction functions and the parameters used for this network can be found respectively in the Appendices D and E.

**Figure 3:**
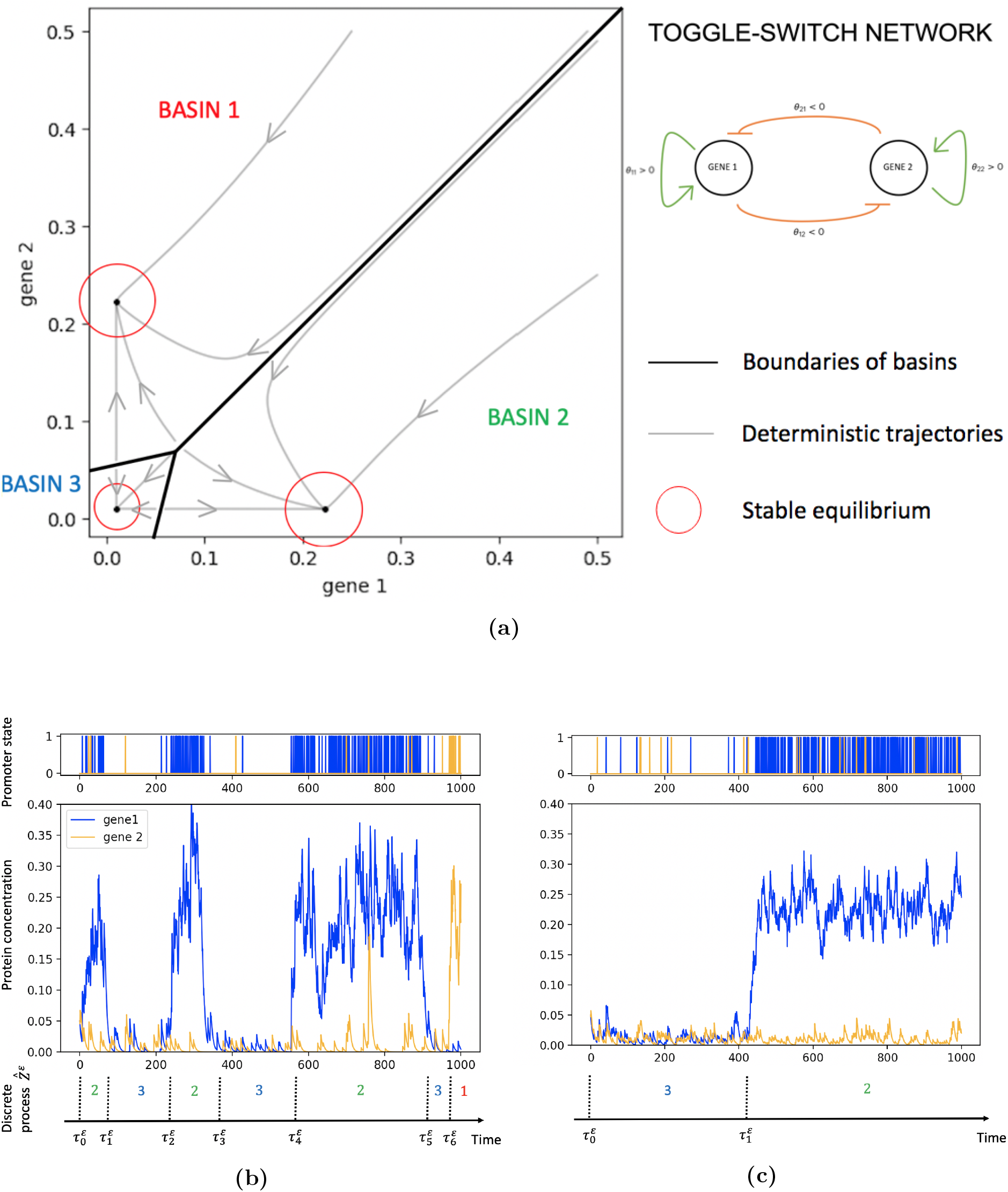
3a: Phase portrait of the deterministic approximation for a symmetric toggle-switch with strong inhibition: two genes which activate themselves and inhibit each other. 3b: Example of a stochastic trajectory generated by the toggle-switch, for *ε* = 1/7. 3c: Example of a stochastic trajectory generated by the toggle-switch, for *ε* = 1/30.

### 2.2 Metastability

When the parameter *ε* is small, transitions from one basin of attraction to another are rare events: in fact the mean escape time from each basin is much larger than the time required to reach a local equilibrium (quasi-stationary state) in the basin.

Adopting the paradigm of metastability mentioned in the introduction, we identify each cell type to a basin of attraction associated to a stable equilibrium of the deterministic system (4). In this point of view, a cell type corresponds to a metastable sub-region of the gene expression space. It also corresponds to the notion of macrostate used in the theory of Markov State Models which has been recently applied to a discrete cell differentiation model in [37]. Provided we can describe accurately the rates of transition between the basins on the long run, the process can then be coarsely reduced to a new discrete process on the cell types.

More precisely, let *m* be the number of stable equilibrium points of the system (4), called attractors. We denote *Z* the set of the *m* basins of attraction associated to these *m* attractors, that we order arbitrarily: *Z* = {*Z*_1_, · · ·, *Z*_*m*_}. The attractors are denoted by 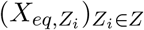. Each attractor is associated to a unique basin of attraction. By assumption, the closure of these *m* basins, 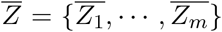, covers the gene expression space Ω. To obtain an explicit characterization of the metastable behavior, we are going to build a discrete process 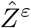, with values in *Z*. From a random path 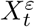 of the PDMP system such that 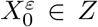, we define a discrete process 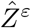 describing the cell types:

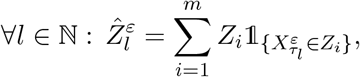

where 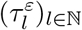 is a sequence of stopping times defined by: 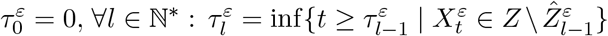. Note that 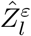 are the successive metastable states, and that 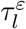 are the successive times of transition between them. From the convergence of any random path to a solution of the deterministic system (4), that we mentioned in Section 2.1, we know that for every basin *Z*_*i*_ such that 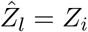, whatever is the point on the boundary of *Z*_*i*_ which has been first attained, 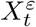 reaches any small neighborhood of the attractor 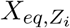 of *Z*_*i*_ before leaving the basin, with probability converging to 1 as *ε* → 0. In addition, for any basin *Z*_*j*_, the probability 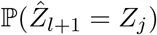 is asymptotically independent of the value of 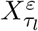, and is then asymptotically determined by the value of 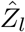. In other words, 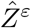 converges to a Markov chain when *ε* → 0. We refer to [38] to go further in the analysis of the coupling between the processes 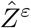 and *X*^*ε*^ for general Markov processes.

For small *ε*, it is natural to approximate the distribution of the exit time from a basin by an exponential distribution. The *in silico* distribution represented in Figure 4 suggests that this assumption seems accurate for the toggle-switch network, even for a realistic value of *ε*. Note, however, that the exponential approximation slightly overestimates the probability that the exit times are small.

**Figure 4:**
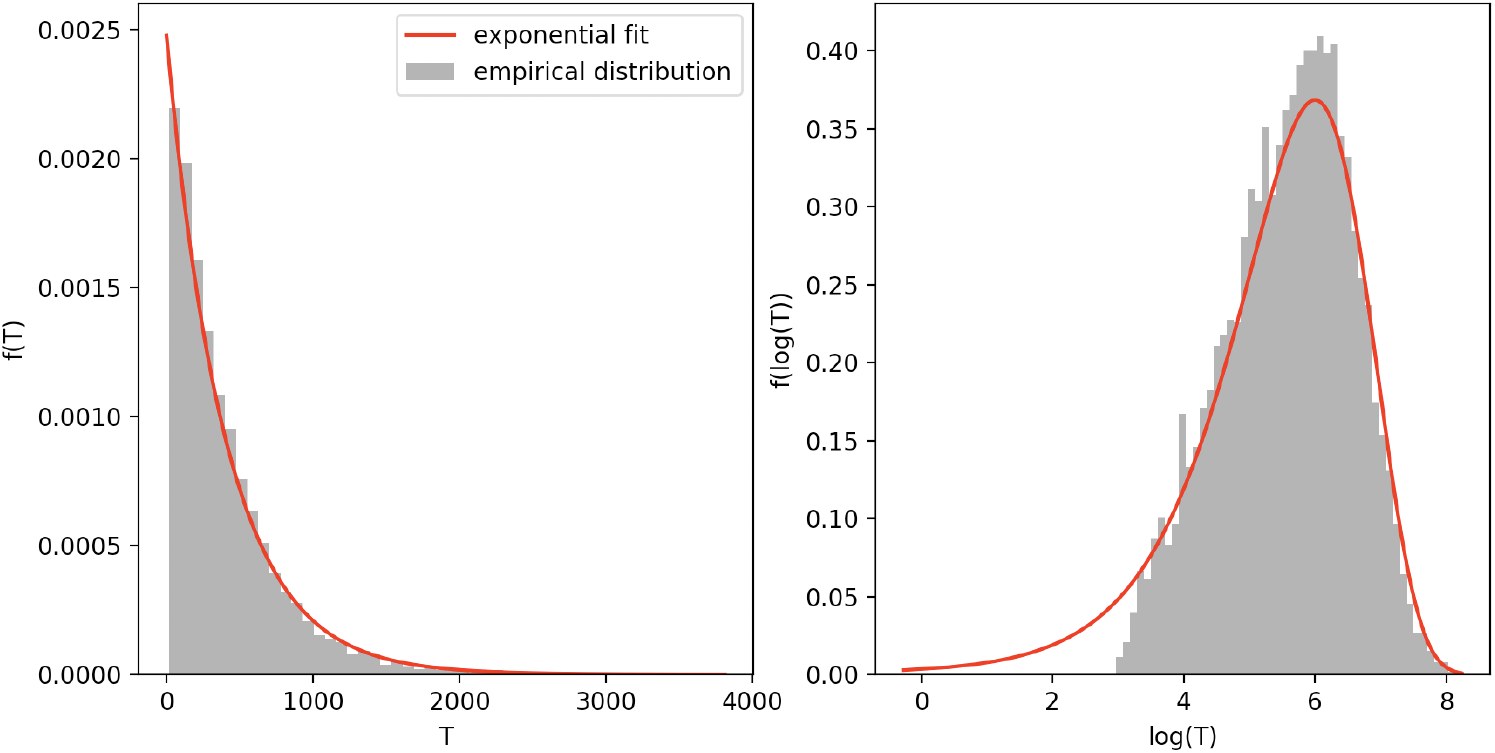
Comparison between the distribution of the exit time from a basin, obtained with a Monte-Carlo method, and the exponential law with appropriate expected value, for *ε* = 1/7. We represent the two densities in normal scale (on the left-hand side) and in logarithmic scale (on the right-hand side) to observe that the exponential law overestimates the probability that the exit times are small.

To completely characterize the coarse-grained resulting process, it remains to compute the transition rates 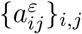 of the time-continuous Markov chain on the basins, that we define for all pair of basins (*Z*_*i*_, *Z*_*j*_) ∈ *Z*^2^, *i* ≠ *j*, by:

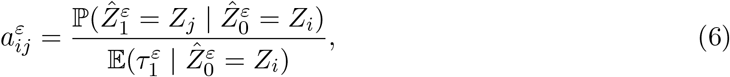

where 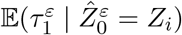 is called the Mean First Exit Time of exit from *Z*_*i*_. This Markov process with discrete state space *Z* represents accurately, when *ε* is small enough, the main dynamics of the metastable system in the weak noise limit [39]. This reduced model is fully described by *m*^2^ transition rates: when the number of genes *n* is large, it is significantly smaller than the *n*^2^ parameters characterizing the GRN model (see Appendix D).

This collection of transition rates are characterized by rare events: when *ε* ≪ 1 or when the number of genes is large, it would be too expensive to compute them with a crude Monte-Carlo method. We are then going to present a method for approximating these transition rates from probabilities of some rare events. We will detail afterwards how these probabilities can be computed either by an efficient numerical method or an analytical approximation.

## 3 Computing the transition rates

### 3.1 Transition rates from probabilities of rare events

In this Section, we approximate each transition rate between any pair of basins (*Z*_*i*_,*Z*_*j*_), *j* ≠ *i* in terms of the probability that a random path realizes a certain rare event in the weak noise limit.

Let us consider two small parameters *r, R* such that 0 < *r* < *R*. We denote 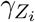 the *r*-neighborhood of 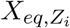, and 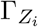 its *R*-neighborhood. For a random path 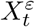 of the PDMP system starting in 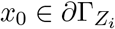, we denote the probability of reaching a basin *Z*_*j*_, *j* ≠ *i* before any other basin *Z_k_*, *k* ≠ *i, j*, and before entering in 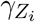:

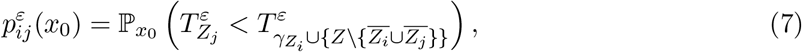

where 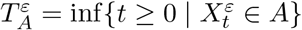 is the hitting time of a set *A* ⊂ Ω.

The method developed in [40] aims to show how it is possible, from the knowledge of the probability (7), to approximate the transition rate 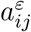 presented in (6). Briefly, it consists in cutting a transition path into two pieces, a piece going from 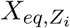 to 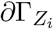 and another reaching *Z*_*j*_ from 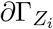: the transition rates *a*_*ij*_ can be then approximated by the reverse of the mean number of attempts of reaching *Z*_*j*_ from 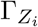 before entering in 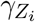, which is close to the inverse of the rare event probability given by (7) when 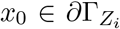, multiplied by the average time of each excursion, that we denote 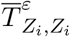. We obtain:

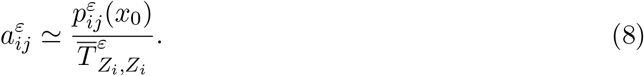

It is worth noticing that to be rigorous, this method need to redefine the neighborhoods 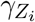 and 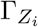 by substituting to the squared euclidean distance a new function based on the probability of reaching the (unknown) boundary: 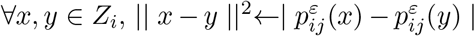. The details are provided in Appendix F.

We observe that the average time 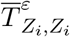 can be easily computed by a crude Monte-Carlo method: indeed, the trajectories entering in 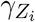 are not rare. It thus only remains to explain the last ingredient for the approximation of the transition rates, which is how to estimate the probabilities of the form (7).

### 3.2 Computing probabilities of the form (7)

A powerful method for computing probabilities of rare events like (7), is given by splitting algorithms. We decide to adapt the Adaptative Multilevel Splitting Algorithm (AMS) described in [41] to the PDMP system: all the details concerning this algorithm can be found in Appendix G. In Section 5, we will verify that the probabilities given by the AMS algorithm are consistent with the ones obtained by a crude Monte-Carlo method for the toggle-switch network.

However, estimating the probability (7) becomes hard when both the number of genes of interest increases and *ε* decreases. Indeed, the AMS algorithm allows to compute probabilities much smaller than the ones we expect for biologically relevant parameters (*ε* ≈ 0.1), but the needed number of samples grows at least with a polynomial factor in *ε*^−1^. If the number of genes considered is large, these simulations can make the algorithm impossible to run in a reasonable time. A precise analysis of the scalability of this method for the PDMP system is beyond the scope of this article, but we have been able to get consistent results on a laptop in time less than one hour for a network of 5 genes, with *ε* > 1/15. The resulting probabilities were of order 5.10^−3^.

In order to overcome this problem, we are now going to develop an analytical approach for approximating these probabilities, by means of optimal trajectories exiting each basin of attraction. The later can be computed within the context of Large deviations. As we will see in Section 5, this approach is complementary to the AMS algorithm, and consistent with it.

#### 3.2.1 Large deviations setting

In this section, we derive a variational principle for approximating the transition probability (7) introduced in Section 3.1. A powerful methodology for rigorously deriving a variational principle for optimal paths is Large deviations theory. It has been developed extensively within the context of Stochastic Differential Equations (SDE) [39][42]. For the sake of simplicity, we present here only an heuristic version. There exists a Large deviations principle (LDP) for a stochastic process with value in Ω if for all *x*_0_ ∈ Ω there exists a lower semi-continuous function defined on the set of continuous trajectories from [0, *T*] to Ω, *J*_*T*_ : *C*_0*T*_ (ℝ^*n*^) → [0, ∞], such that for all set of trajectories *A* ⊂ *C*_0*T*_ (ℝ^*n*^):

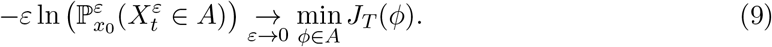

The function *J*_*T*_ is called the rate function of the process in [0, *T*], and the quantity *J*_*T*_ (*ϕ*) is called the cost of the trajectory *ϕ* over [0, *T*].

The particular application of this theory to stochastic hybrid systems has been developed in detail in [43] and [35]. We now present consequences of results developed in [29].

##### Definition 1.

*The Hamiltonian is the function H*: Ω × ℝ^*n*^ ↦ ℝ, *such that for all* (*x, p*) ∈ Ω × ℝ^*n*^, *H*(*x, p*) *is the unique eigenvalue associated to a nonnegative right-eigenvector* 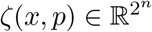, *(which is unique up to a normalization), of the following spectral problem:*

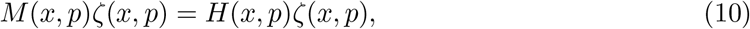

*where the matrix* 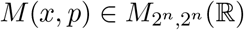 *is defined by:*

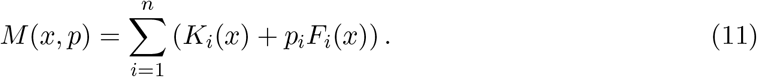

We remark that the matrix *M* (*x, p*) has off-diagonal nonnegative coefficients. Moreover, the positivity of the functions *k*_*on,i*_ makes *M* irreducible (the matrix allows transition between any pair 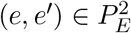 after at most *n* steps). Thereby, the Perron Frobenius Theorem may be applied, and it justifies the existence and uniqueness of *H*(*x, p*). Moreover, from a general property of Perron eigenvalues when the variable *p* appears only on the diagonal of the matrix, *H* is known to be convex [44].

The following result is a direct consequence of theoretical results of [29] applied to the PDMP system (1).

##### Theorem 1.

*Let us denote 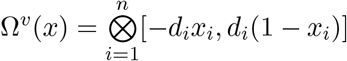 and* 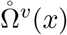 *its interior.*

*The Fenchel-Legendre transform of the Hamiltonian H is well-defined and satisfies:* ∀_*x*_ ∈ Ω, 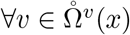,

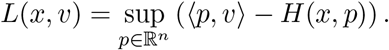

*Moreover, the PDMP system* (1) *satisfies a LDP, and the associated rate function has the form of a classical action: the cost of any piecewise differentiable trajectory ϕ_t_ in C*_0*T*_ (ℝ^*n*^) *satisfying for all t* ∈ [0, *T*), 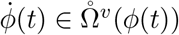, *can be expressed as*

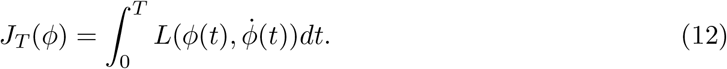

The function *L* is called the Lagrangian of the PDMP system.

We are now going to show how in certain cases, trajectories which minimize the quantity (12) between two sets in Ω can be defined with the help of solutions of an Hamilton-Jacobi equation with Hamiltonian *H*.

#### 3.2.2 WKB approximation and Hamilton-Jacobi equation

The Hamiltonian defined in (10) also appears in the WKB (Wentzell, Kramer, Brillouin) approximation of the master equation [34], [29]. This approximation consists in injecting in the master equation (2) of the PDMP system, a solution of the form:

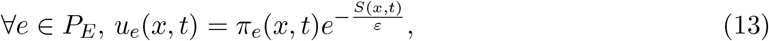

where 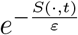 then denotes the marginal distribution on proteins of the distribution *u* at time *t*, and *π*_*e*_(*x, t*) is a probability vector denoting the conditional distribution of the promoters knowing that proteins are fixed to *X* = *x* at *t*. The expression (13) is justified under the assumption that the density *u*_*e*_ is positive at all times.

Under the regularity assumptions *S* ∈ *C*^1^(Ω × ℝ^+^, ℝ) and 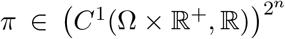, we can perform a Taylor expansion in *ε* of the functions *S* and *π*_*e*_, for any *e* ∈ *P*_*E*_, and keeping in the resulting master equation only the leading order terms in *ε*, that we denote *S*_0_ and 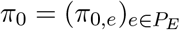, we obtain:

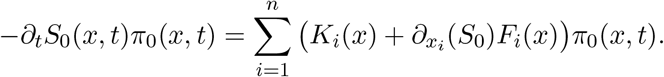

Identifying the vectors *π*_0_ and ∇_*x*_*S*_0_ with the variables *ζ* and *p* in equation (10), we obtain that *S*_0_ is solution of an Hamilton-Jacobi equation:

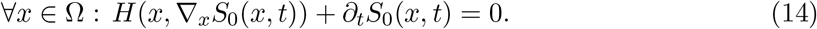

More precisely, if at any time *t*, the marginal distribution on proteins of the PDMP process, denoted *u*(·, *t*), follows a LDP and if its rate function, denoted *V* (·, *t*), is differentiable on Ω, then the function *H*(·, ∇_*x*_*V* (·, *t*)) appears as the time derivative of *V* (·, *t*) at *t*. Moreover, the WKB method presented above shows that the rate function *V* (·, *t*) is identified for any time *t* with the leading order approximation in *ε* of the function *S*(·, *t*) = −*ε* log(*u*(·, *t*)). Note that (13) is also reminiscent of the Gibbs distribution associated with a potential *S*. Some details about the interpretation of this equation and its link with the quasistationary approximation can be found in [34].

Next, we consider a function *V*, solution of the Hamilton-Jacobi equation (14), that we assume being of class *C*^1^(Ω × ℝ^+^, ℝ) for the sake of simplicity. Then, for any piecewise differentiable trajectory *ϕ*_*t*_ ∈ *C*_0*T*_ (Ω) such that 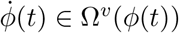 for all *t* ∈ [0, *T*), one has, by definition of the Fenchel-Legendre transform:

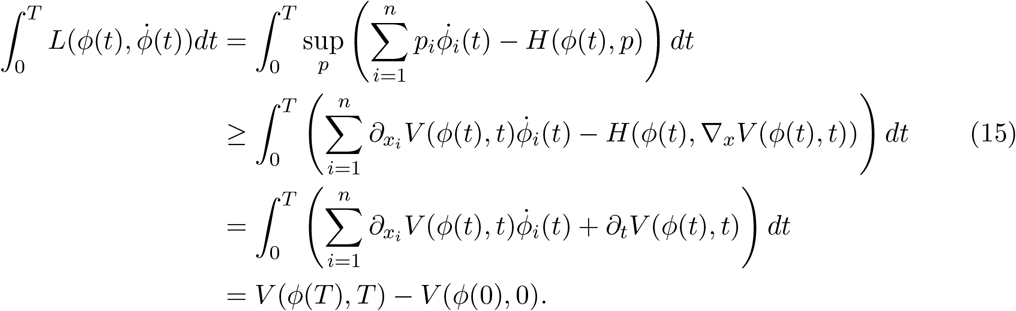

Moreover, when *H* is strictly convex in *p*, we have:

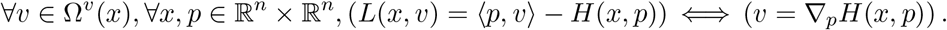

Then, the equality in (15) is exactly reached at any time for trajectories *ϕ*_*t*_ such that for all *t* ∈ [0, *T*), *i* = 1, · · ·, *n*:

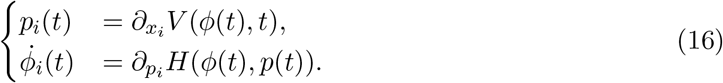

#### 3.2.3 General method for computing probabilities of the form (7)

We now detail the link existing between the regular solutions *V* of the Hamilton-Jacobi equation (14) and the probabilities of the form (7). For this, we introduce the notion of quasipotential.

##### Definition 2.

*Denoting* 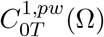 *the set of piecewise differentiable trajectories in C*_0*T*_ (Ω)*, we define the quasipotential as follows: for two sets A, B* ⊂ Ω *and a set R* ⊂ Ω \ (*A* ∪ *B*),

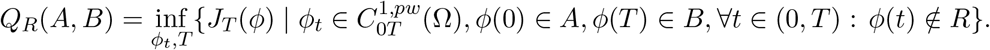

*We call a trajectory* 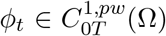 *an optimal trajectory between the two subsets A, B* ⊂ Ω *in* Ω \ *R, if it reaches the previous infimum.*

For the sake of simplicity, if *R* = ∅, we will write *Q*_*R*_(*A, B*) = *Q*(*A, B*).

With these notations, the LDP principle allows to approximate for any basin *Z*_*j*_, *i* ≠ *j*, the probability 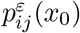 defined in (7), which is the probability of reaching *Z*_*j*_, from a point 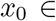 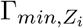, before 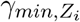, by the expression:

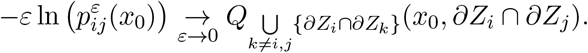

A direct consequence of the inequality (15), in the case where equality is reached, is that a regular solution *V* of the Hamilton-Jacobi equation (14) defines trajectories for which the cost (12) is minimal between any pair of its points. Moreover, if *V* is a stationary solution of (14), the cost of such trajectories does not depend on time: these trajectories are then optimal between any pair of its points among every trajectory in any time. We immediately deduce the following lemma:

#### Lemma 1.

*For a stationary solution V* ∈ *C*^1^(Ω, ℝ) *of* (14) *and for all T* > 0*, any trajectory* 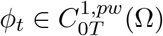 *satisfying the system* (16) *associated to V is optimal in* Ω *between ϕ*(0) *and ϕ*(*T*)*, and we have:*

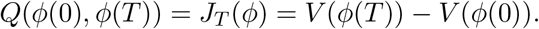

Thus, for approximating the probability of interest (7), between any pair of basin (*Z*_*i*_, *Z*_*j*_), we are going to build a trajectory 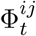, which verifies the system (16) associated to a stationary solution *V* of (14), with 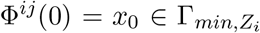, and which reaches in a time *T* a point *x* ∈ *∂Z*_*i*_ ∩ *∂Z*_*j*_ such that 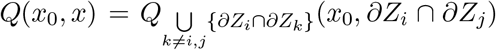. For such trajectory, from Lemma 1, we could then approximate the probability (7) by the formula

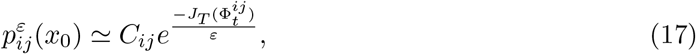

where *C*_*ij*_ is an appropriate prefactor. Unfortunately, if there exists an explicit expression of *C*_*ij*_ in the one-dimensional case [34], and that an approximation has been built for multi-dimensional SDE model [45], they are intractable or not applicable in our case. In general, the prefactor does not depend on *ε* [46]. In that case − ln 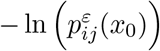 is asymptotically an affine function of *ε*^−1^, the slope of which is 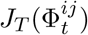 and the initial value − ln(*C*_*ij*_). Then, the strategy we propose simply consists in approximating the prefactor by comparison between the probabilities given by the AMS algorithm and the Large deviations approximation (17) for a fixed *ε* (large enough to be numerically computed.)

To conclude, for every pair of basins (*Z*_*i*_, *Z*_*j*_), *i* ≠ *j*, one of the most efficient methods for computing the probability (7) is to use the AMS algorithm. When the dimension is large, and for values of *ε* which are too small for this algorithm to be efficiently run, we can use the LDP approximation (17), provided that the corresponding optimal trajectories 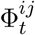 can be explicitly found. The latter condition is studied in the next sections. The AMS algorithm is then still employed to approximate the prefactor, which is done using intermediate values of *ε* by the regression procedure mentioned above.

## 4 Analytical approximation of probabilities of the form (7) for the PDMP system

### 4.1 Expressions of the Hamiltonian and the Lagrangian

In this section, we identify the Perron eigenvalue *H*(*x, p*) of the spectral problem (10), and prove that its Fenchel-Legendre transform *L* with respect to the variable *p* is well defined on ℝ^*n*^. We then obtain the explicit form of the Hamiltonian and the Lagrangian associated to the LDP for the PDMP system (1).

#### Theorem 2.

*For all n in* ℕ\**, the Hamiltonian is expressed as follows: for all* (*x, p*) ∈ Ω × ℝ^*n*^, *the unique solution of the spectral problem* (10) *(with nonnegative eigenvector) is:*

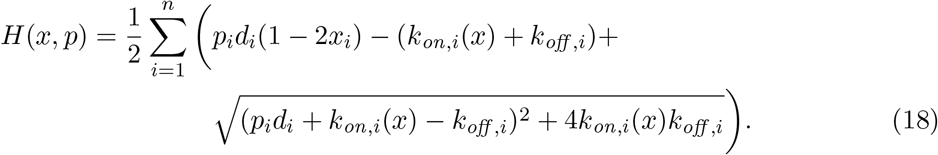

*Moreover, the function H is strictly convex with respect to p.*

#### Theorem 3.

*The Lagrangian is expressed as follows: for all* (*x, v*) ∈ Ω × ℝ^*n*^, *one has*:

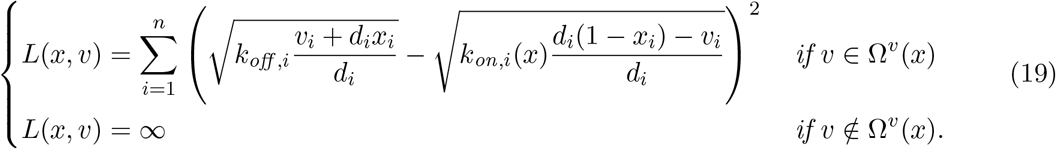

*In addition, for all x* ∈ Ω*, L*(*x, v*) = 0 *if and only if for all i* = 1, · · ·, *n:*

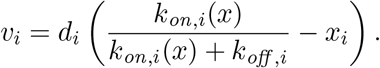

As detailed in Appendix C.2, we remark that the Lagrangian of the PDMP process defined in (19) is not equal to the Lagrangian of the diffusion approximation defined in Appendix C.1, which is:

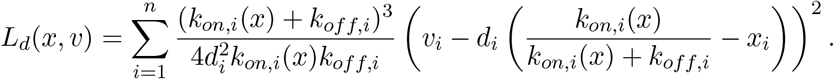

More precisely, the Lagrangian of the diffusion approximation is a second order approximation of the Taylor expansion of the Lagrangian of the PDMP system around the velocity field associated to the deterministic limit system (4). Observe that the Lagrangian of the diffusion approximation is a quadratic mapping in *v*, which is expected since the diffusion approximation is described by a Gaussian process. On the contrary, the Lagrangian *L* given by (19) is not quadratic in *v*. As it had been shown in [47] for Fast-Slow systems, this highlights the fact that the way rare events arise for the PDMP system is fundamentally different from the way they would arise if the dynamics of proteins was approximated by an SDE.

*Proof of Theorem 2.* Defining the 2 × 2 matrix

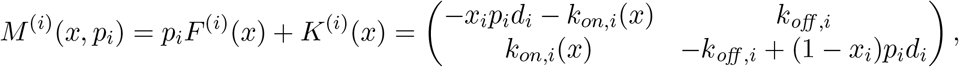

the Perron eigenproblem associated to *M* ^(*i*)^

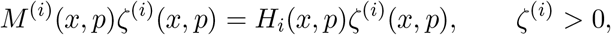

implies immediately that

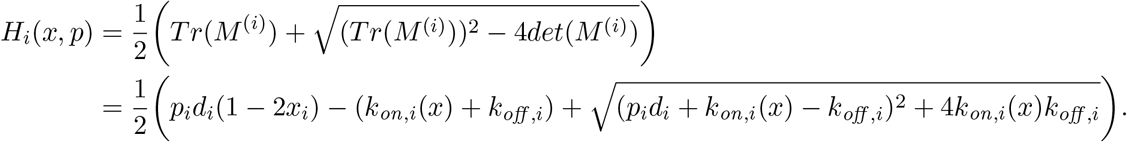

If we impose the constraint 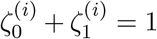, *i.e* that there exists for all *x*, *p*_*i*_, *α*_*p,i*_(*x*) ∈ (0, 1) such that 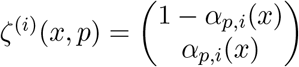, we obtain the following equation:

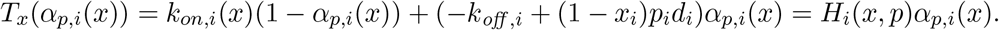

Since *T*_*x*_(0) = −*k*_*on,i*_(*x*) and *T*_*x*_(1) = *k*_*off,i*_, *T*_*x*_ has one and only one root in (0, 1). After a quick computation, one gets for all *x, p* ∈ Ω × ℝ^*n*^:

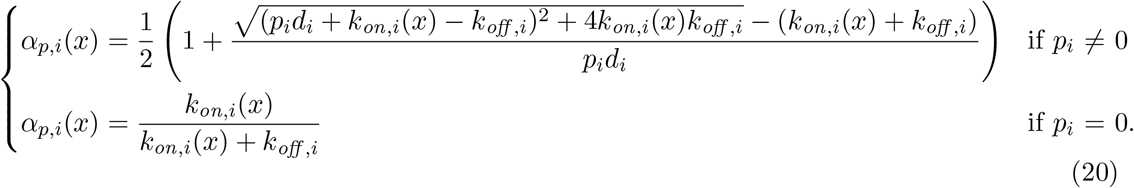

Considering the tensorial structure of the problem, denoting *M*_*i*_(*x, p*) = *p*_*i*_*F*_*i*_(*x*) + *K*_*i*_(*x*) (the tensorial version, see Section 1), we have by definition of *M* (11):

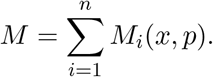

For 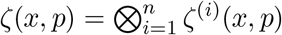, we obtain:

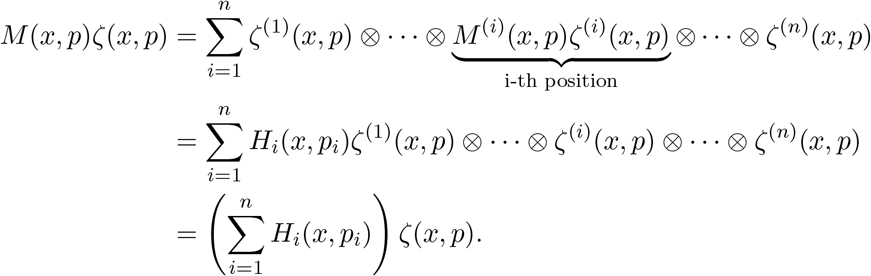

Since *ζ* > 0, one obtains the expression (18) for the Hamiltonian:

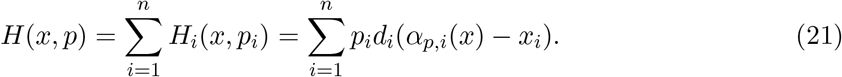

We verify that *H* is strongly convex with respect to *p*, which follows from the following computation: for all *i, j* = 1, · · ·, *n,*

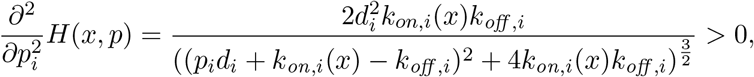

and the cross-derivatives are clearly 0. This concludes the proof of Theorem 2.

*Proof of Theorem 3.* The objective is to compute the Fenchel-Legendre transform of the Hamiltonian *H* given by (18) in Theorem 2.

For all *x* ∈ Ω and for all *v*_*i*_ ∈ ℝ, the function *g*: *P*_*i*_ ↦ *p*_*i*_*v*_*i*_ − *H*_*i*_(*x*, *p*_*i*_) is concave. An asymptotic expansion (when *p*_*i*_ → ±∞) gives:

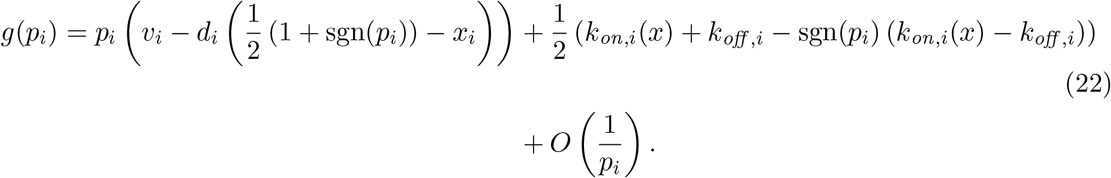

Let us study three cases. If *v*_*i*_ ∈ (−*d*_*i*_*x*_*i*_, *d*_*i*_(1 − *x*_*i*_)), *g* goes to −∞ when *p*_*i*_ → ±∞: thus *g* reaches a unique maximum in ℝ. At the boundary *v*_*i*_ = −*d*_*i*_*x*_*i*_ (resp. *v*_*i*_ = *d*_*i*_(1 − *x*_*i*_)), *g* goes to −∞ as *P*_*i*_ goes to +∞ (resp. −∞) and converges to *k*_*on,i*_(*x*) (resp. *k*_*off,i*_) as *P*_*i*_ goes to −∞ (resp. +∞): then *g* is upper bounded and the sup is well defined. If *v*_*i*_ ∉ [−*d*_*i*_*x*_*i*_, *d*_*i*_(1 − *x*_*i*_)], *g*(*P*_*i*_) goes to +∞ when either *P*_*i*_ → −∞ of *P*_*i*_ → +∞, thus *g* is not bounded from above. As a consequence, 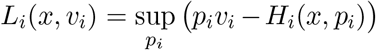 is finite if and only if *v*_*i*_ ∈ [−*d*_*i*_*x*_*i*_,*d*_*i*_(1 − *x*_*i*_)].

The Fenchel-Legendre transform of *H* is then given as follows: for all *x* ∈ Ω and *v* ∈ ℝ^*n*^

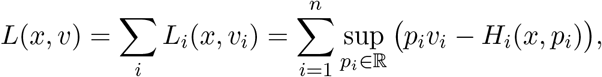

and *L*(*x, v*) is finite for every *v* ∈ Ω^*v*^(*x*). To find an expression for *L*(*x, v*), we have to find for all *i* = 1, · · ·, *n* the unique solution *p*_*v,i*_(*x*) of the invertible equation: 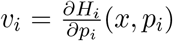. Developing the term on the right-hand side, we obtain:

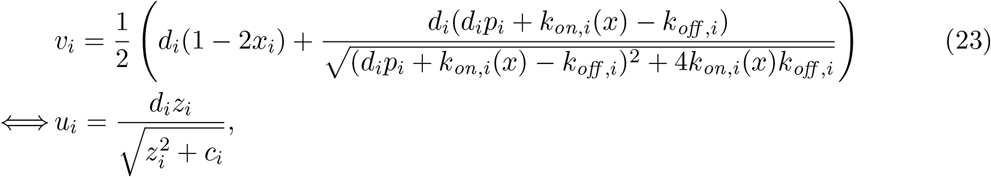

where *u*_*i*_ = 2(*v*_*i*_ + *d*_*i*_*x*_*i*_) − *d*_*i*_, *c*_*i*_ = 4*k*_*on,i*_(*x*)*k*_*off,i*_ > 0, *z*_*i*_ = *d*_*i*_*p*_*i*_ + *k*_*on,i*_(*x*) − *k*_*off,i*_.

When *v*_*i*_ ∈ (−*d*_*i*_*x*_*i*_, *d*_*i*_(1 − *x*_*i*_)), we have *u*_*i*_ ∈ (−*d*_*i*_, *d*_*i*_). Thus, we obtain

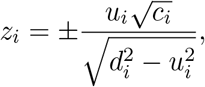

and as *Z*_*i*_ and *u*_*i*_ must have the same sign, we can conclude for every *v*_*i*_ ∈ (−*d*_*i*_*x*_*i*_, *d*_*i*_(1 − *x*_*i*_)):

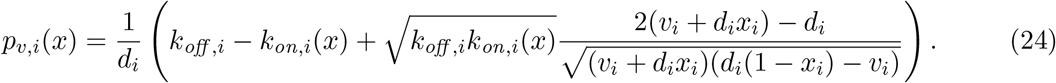

Injecting this formula in the expression of the Fenchel-Legendre transform, we obtain after straightforward computations:

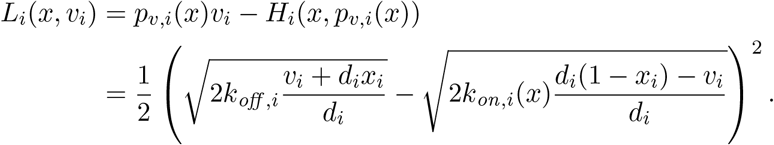

We finally obtain the expression when *v* ∈ Ω^*v*^(*x*):

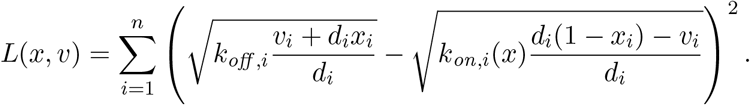

Finally, if *v* ∉ Ω^*v*^(*x*), *i.e.* if there exists *i* such that *v*_*i*_ ∉ [−*d*_*i*_*x*_*i*_, *d*_*i*_(1 − *x*_*i*_)], then *L*(*x, v*) = *L*_*i*_(*x*_*i*_, *v*_*i*_) = ∞. As expected, the Lagrangian is always nonnegative. In addition, it is immediate to check that *L*(*x, v*) = 0 if and only if the velocity field *v* is the drift of the deterministic trajectories, defined by the system (4).

### 4.2 Stationary Hamilton-Jacobi equation

We justified in Section 3.2.3 that the stationary solutions of the Hamilton-Jacobi equation (14) are central for finding an analytical approximation of the transition rates described in Section 3.1. Thus, we are going to study the existence and uniqueness (under some conditions) of functions *V* ∈ *C*^1^(Ω, ℝ) such that for all *x* ∈ Ω:

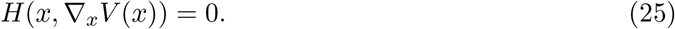

Recalling that from (21), 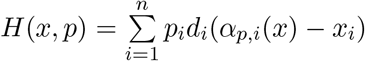, we construct two classes of solutions *V*, such that for all 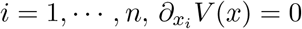 or 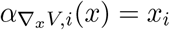.

The first class of solutions contains all the constant functions on Ω. From the expression (20). The second class contains all functions *V* such that for all *x* ∈ Ω:

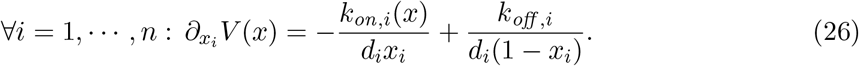

In particular, we show in Appendix D that the condition (26) holds for the toggle-switch network described in Appendix E and studied in Section 5.

We will see in the next section that the class of constant solutions are associated to the deterministic system (4), which are the trajectories of convergence within the basins. We describe in Section 4.3.2 a more general class of solutions than (26), which defines the optimal trajectories of exit from the basins of attraction of the deterministic system.

### 4.3 Optimal trajectories

In the sequel we study some properties of the optimal trajectories associated to the two classes of solutions of the stationary Hamilton-Jacobi equation (25) introduced above.

#### 4.3.1 Deterministic limit and relaxation trajectories

From Lemma 1, for every constant function *V* (·) = *C* on Ω, the associated collection of paths *ϕ*_*t*_ satisfying the system (16) is optimal in Ω between any pair of its points. Replacing *p* = ∇_*x*_*V* = 0 in (16), we find that these trajectories verify at any time *t* > 0 the deterministic limit system (4):

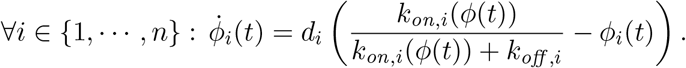

Moreover, for every trajectory *ϕ*_*t*_ solution of this system, we have for any *T* > 0:

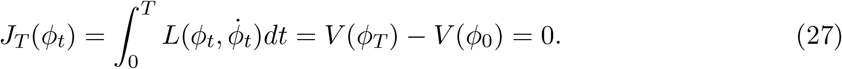

We call such trajectories the *relaxation trajectories*, as they characterize the optimal path of convergence within every basin. From Theorem 3, these relaxation trajectories are the only zero-cost admissible trajectories.

#### 4.3.2 Quasipotential and fluctuation trajectories

We now characterize the optimal trajectories of exit from the basins. We are going to show that the condition (C) defined below is sufficient for a solution *V* ∈ *C*^1^(Ω, ℝ) of the equation (25) to define optimal trajectories realizing the rare events described by the probabilities (7).

##### Definition 3.

*We define the following condition on a function V* ∈ *C*^1^(Ω, ℝ): *(C) The set* {*x* ∈ Ω | ∇_*x*_*V* (*x*) = 0} *is reduced to isolated points.*

The results presented below in Theorem 4 are mainly an adaptation of Theorem 3.1, Chapter 4, in [39]. In this first Theorem, we state some properties of solutions *V* ∈ *C*^1^(Ω, ℝ) of (25) satisfying the condition (C):

##### Theorem 4.

*Let V* ∈ *C*^1^(Ω, ℝ) *be a solution of* (25).

i. *For any optimal trajectory ϕ_t_ satisfying the system* (16) *associated to V, for any time t we have the equivalence:*

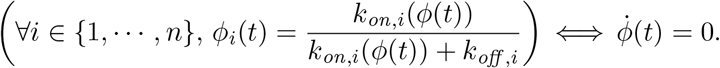
ii. *The condition (C) implies that the gradient of V vanishes only on the stationary points of the system* (4).
iii. *If V satisfies (C), then V is strictly increasing on any trajectory which solves the system* (16)*, such that the initial condition is not an equilibrium point of the system* (4)*. Moreover, for any basin of attraction Z_i_ associated to an attractor 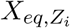, we have:*

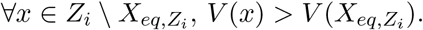
iv. *If V satisfies the condition (C), under the assumption that* 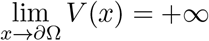, *we have the formula:*

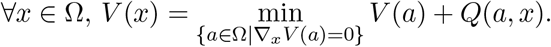
v. *Let us consider V,* 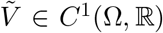 *two solutions of* (25) *satisfying the condition (C). The stable equilibria of the system defined for every time t by 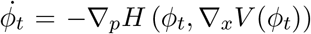 *are exactly the attractor of the deterministic system* (4) 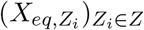. We denote* 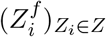 *the basins of attraction which are associated to these equilibria: at least on 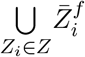, the relation* 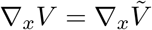 *is satisfied.*

*Moreover, under the assumptions 1. that* 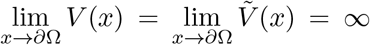, *and 2. that between any pair of basins* 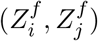, *we can build a serie of basins* 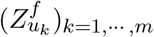 *such that u*_0_ = 1, *u*_*m*_ = *j and for all k < m*, 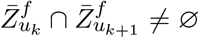, *then V and* 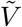 *are equal in* Ω *up to a constant.*

Note that the point (iii) makes these solutions consistent with the interpretation of the function *V* as the rate function associated to the stationary distribution of the PDMP system, presented in Section 3.2.2. Indeed, as every random path converges in probability when *ε* → 0 to the solutions of the deterministic system (4) [35], the rate function has to be minimal on the attractors of this system, which then corresponds to the points of maximum likelihood at the steady state. It should also converge to +∞ on *∂*Ω, as the cost of reaching any point of the boundary is infinite (see Corollary 2, in the proof of Theorem 5). However, we see in (v) that the uniqueness, up to a constant, needs an additional condition on the connection between basins which remains not clear for us at this stage, and which will be the subject of future works.

If *V* ∈ *C*^1^(Ω, ℝ) is a solution of (25) saitsfying (C), we call a trajectory solution of the system (16) associated to *V* a *fluctuation trajectory*.

We observe that any function satisfying the relation (26) belongs to this class of solutions of (25), and then that in particular, such *C*^1^ function exists for the toggle-switch network. In that special case, we can explicitly describe all the fluctuation trajectories: for any time *t*, replacing 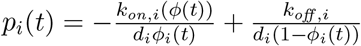 in the system (16), we obtain

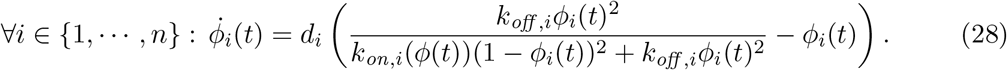

In the second theorem (Theorem 5), we justify that the fluctuation trajectories are the optimal trajectories of exit:

##### Theorem 5.

*Let us assume that there exists a function V* ∈ *C*^1^(Ω, ℝ) *which is solution of* (25) *and satisfies the condition (C). For any basin Z_i_* ∈ *Z, there exists at least one basin Z_j_, j* ≠ *i, such that there exists a couple* (*x*_0_, ϕ_t_), *where* 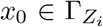 *and ϕ_t_ is a fluctuation trajectory, and such that ϕ*(0) = *x*_0_, 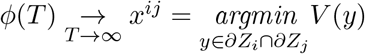.

*Let us denote* 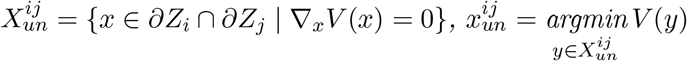 *and* 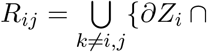 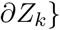. *Under the following assumption y*∈*X^ij^*

*(A) any relaxation trajectory starting in ∂Z_i_* ∩ *∂Z_j_ stays in ∂Z_i_* ∩ *∂Z*_*j*_, *we have* 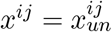 *and:*

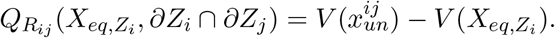

In particular, if there exists a fluctuation trajectory between any attractor 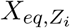 and every saddle points of the deterministic system (16) on the boundary *∂Z*_*i*_, and if the assumption (A) of Theorem 5 is verified for every basin *Z*_*j*_, *j* ≠ *i*, the function *V* allows to quantify all the optimal costs of transition between the basins. This is generally expected because the attractors are the only stable equilibria for the reverse fluctuations (see the proof of Theorem 4.(v)). The proofs of Theorems 4 and 5 use classical tools from Hamiltonian system theory and are postponed to Appendix H.

When a solution *V* ∈ *C*^1^(Ω, ℝ) satisfying (C) exists, the saddle points of the deterministic system (4) are then generally the bottlenecks of transitions between basins and the function *V* characterizes the energetic barrier between them. The function 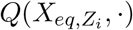 depends on the basin *Z*_*i*_, which is a local property: it explains why the function *V* is generally called the global quasipotential, and 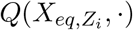 the local quasipotential of the process [48].

The precise analysis of the existence of a regular solution *V* satisfying (C) for a given network is beyond the scope of this article. When it is impossible to find a regular solution, more general arguments developed within the context of Weak KAM Theory can allow to link the viscosity solutions of the Hamilton-Jacobi equation to the optimal trajectories in the gene expression space [49].

### 4.4 Partial conclusion

We have obtained in Theorem 3 the form of the Lagrangian in the variational expression (12) for the rate function *J*_*T*_ associated to the LDP for the PDMP system (1). We have also highlighted the existence and interpretation of two types of optimal trajectories.

The first class consists in relaxation trajectories, which characterize the convergence within the basins. The fact that they are the only trajectories which have zero cost justifies that any random path 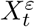 converges in probability to a relaxation trajectory.

When there exists a function *V* ∈ *C*^1^(Ω, ℝ) satisfying (26), the system (28) defines the second class of optimal trajectories, called the fluctuation trajectories. From Theorem 5, for every basin *Z*_*i*_, there exists at least one basin *Z*_*j*_, *j* ≠ *i*, and a trajectory 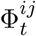 which verifies this system, starts on 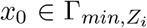 and reaches a point of *x* ∈ *∂Z*_*i*_ ∩ *∂Z*_*j*_ such that *Q*(*x*_0_, *x*) = *Q*(*x*_0_, *∂Z*_*i*_ ∩ *∂Z*_*j*_). This trajectory then realizes the rare event of probability *p*_*ij*_(*x*_0_). Injecting the velocity field defining (28) in the Lagrangian (19), we deduce:

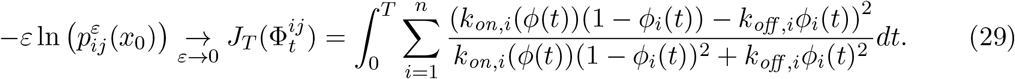

If the assumption (A) of Theorem 5 is verified, this minimum is necessarily reached on a saddle point of *V* on *∂Z*_*i*_ ∩ *∂Z*_*j*_ and in that case, the time *T* must be taken infinite. Then, the formula (29) can be injected in the approximation (17), and the method described in Section 3.2.2 allows to compute the probability of the form (7) for the pair (*Z*_*i*_, *Z*_*j*_).

Moreover, for every basin *Z*_*k*_, *k* ≠ *i*, if the assumption (A) of Theorem 5 is verified and if there exists, for any saddle point 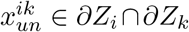, a trajectory satisfying the system (28) which starts at *x*_0_ and reaches 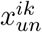 (at *T* → ∞), the formula (29) can also be injected in the approximation (17) for the pair (*Z*_*i*_, *Z*_*k*_), and the method described in Section 3.2.3 allows then to compute the probabilities of the form (7) for any pair of basins (*Z*_*i*_, *Z*_*k*_)_*k*=1,···,*m*_.

## 5 Application to the toggle-switch network

In this section, we consider the class of interaction functions defined in Appendix D for a network with two genes (*n* = 2). This function comes from a chromatin model developed in [28] and is consistent with the classical Hill function characterizing promoters switches. Using results and methods described in the previous sections, we are going to reduce the PDMP system when the GRN is the toggle-switch network described in Appendix E. After defining the attractors of the deterministic system (4), building the optimal fluctuation trajectories between these attractors and the common boundaries of the basins, we will compute the cost of the trajectories and deduce, from the approximation (17), the transition probabilities of the form (7) as a function of *ε*, up to the prefactor. We will compute these probabilities for some *ε* with the AMS algorithm described in Appendix G for obtaining the prefactor. We will then approximate the transition rates characterizing the discrete Markov chain on the cellular types, given by the formula (8), for many values of *ε*. We will finally compare these results to the ones given by a crude Monte-Carlo method.

### 5.1 Computation of the attractors, saddle points and optimal trajectories

First, we compute the stable equilibrium points of the PDMP system (1). The system (5) has no explicit solution. We present a simple method to find them, which consists in sampling a collection of random paths in Ω: the distribution of their final position after a long time approximates the marginal on proteins of the stationary distribution. We use these final positions as starting points for simulating the relaxation trajectories, described by (4), with an ODE solver: each of these relaxation trajectories converges to one of the stable equilibrium points. This method allows to obtain all the stable equilibrium corresponding to sufficiently deep potential wells (see Figure 5). Possible other potential wells can be omitted because they correspond to basins where the process has very low probability of going, and which do not impact significantly the coarse-grained Markov model.

**Figure 5:**
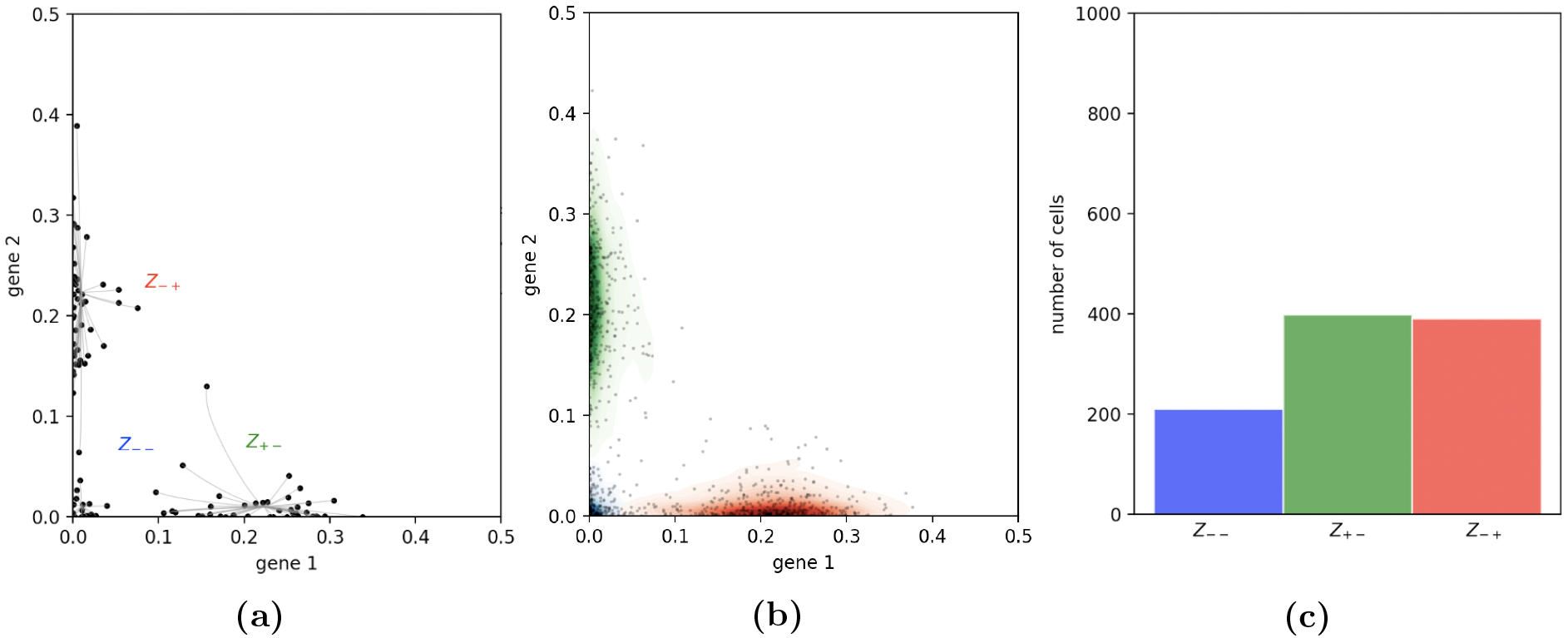
5a: 100 cells are plotted under the stationary distribution. The relaxation trajectories allow to link every cell to its associated attractor. 5b: 1000 cells are plotted under the stationary distribution. They are then classified depending on their attractor, and this figure sketches the kernel density estimation of proteins within each basin. 5c: The ratio of cells that are found within each basin gives an estimation of the stationary distribution on the basins.

Second, we need to characterize the fluctuation trajectories. In Appendix D, we introduced the interaction function and proved that for any symmetric two-dimensional network defined by this function, *i.e* such that for any pair of genes (*i, j*), *θ*_*ij*_ = *θ*_*ji*_ (where *θ* is the matrix characterizing the interactions between genes), there exists a function *V* such that the relation (26) is verified. This is then the case for the toggle-switch network, which is symmetric. We have proved in Section 4.2 that such function *V* solves the Hamilton-Jacobi equation (25), and verifies the condition (C). Thus, the system (28) defines the fluctuation trajectories.

Third, we need to find the saddle points of the system (4). As we know that for any attractor, there exists at least one fluctuation trajectory which starts on the attractor and reaches a saddle point (in an infinite time), a naive approach would consist in simulating many trajectories with different initial positions around every attractors, until reaching many saddle points of the system. This method is called a shooting method and may be very efficient in certain cases. But for the toggle-switch, we observe that the fluctuation trajectories are very unstable: this method does not allow to obtain the saddle points.

We develop a simple algorithm which uses the nonnegative function *l*(·) = *L*(·, ν_v_(·)), which corresponds to the Lagrangian evaluated on the drift ν_v_ of the fluctuation trajectories defined by the system (28). We have:

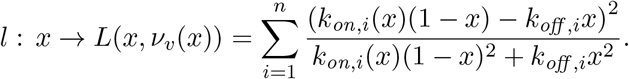

As expected, since ν_v_ cannot be equal to the drift of a relaxation trajectory except on the stationary points of the relaxation trajectories and since the Lagrangian *L*(*x, v*) is equal to 0 if and only if *v* corresponds to the drift of a relaxation trajectory, the function *l* vanishes only on these stationary points. If there exists a saddle point connecting two attractors, this function will then vanish there. The algorithm is described in Appendix I. For the toggle-switch network, it allows to recover all the saddle points of the system (4).

Fourth, we want to compute the optimal trajectories between every attractors and the saddle points on the boundary of its associated basin. Using the reverse of the fluctuation trajectories, for which the attractors of the system (4) are asymptotically stable (see the proof of Theorem 4.(v)), we can successively apply a shooting method around every saddle points. We observe that for the toggle-switch network, for any saddle point at the boundary of two basins, there exists a reverse fluctuation trajectory which converges to the attractors of both basins. For any pair of basins (*Z*_*i*_, *Z*_*j*_), we then obtain the optimal trajectories connecting the attractor 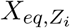 and the saddle points belonging to the common boundary *∂Z*_*i*_ ∩ *∂Z*_*j*_ (see Figure 6).

**Figure 6:**
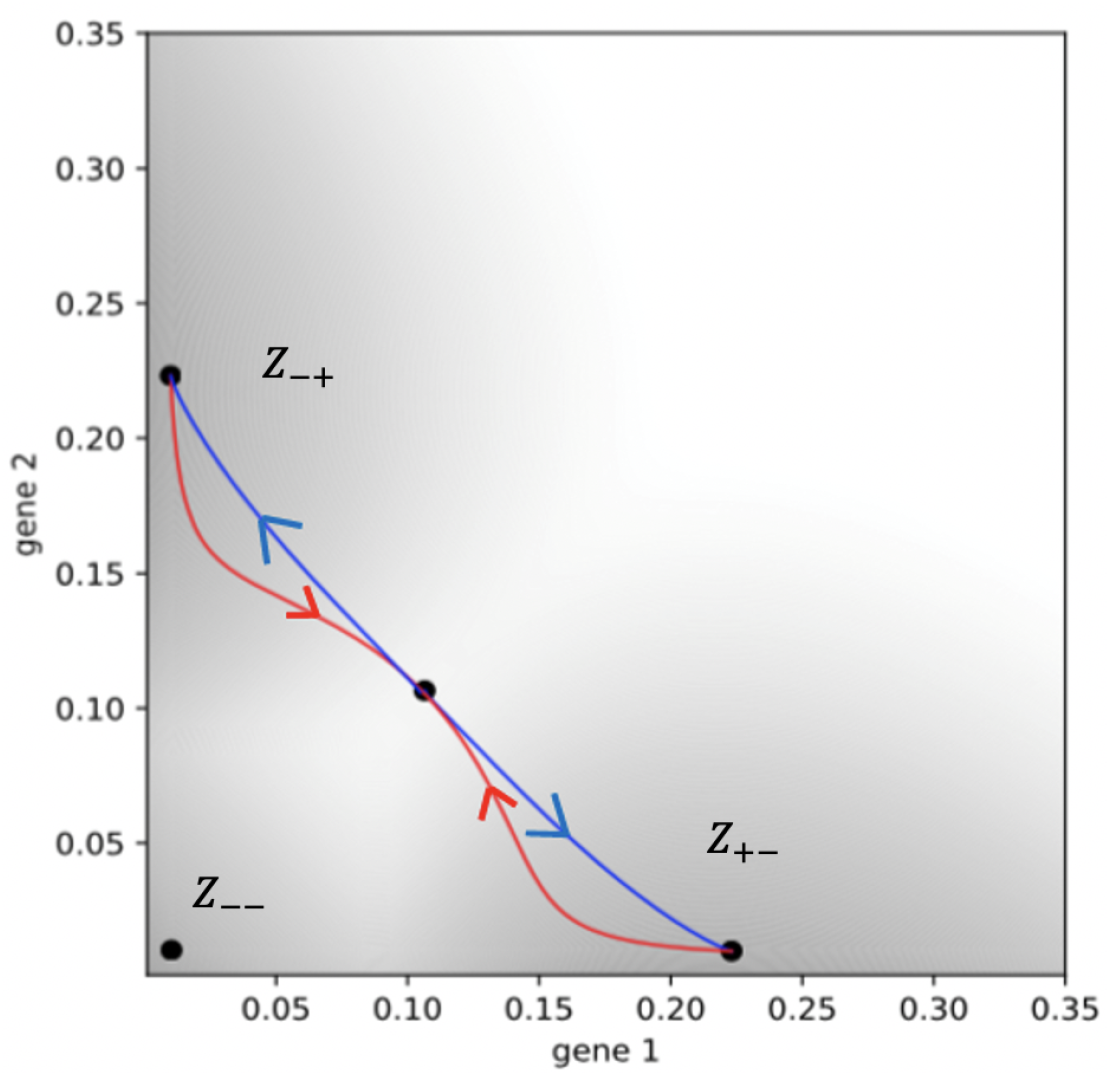
The optimal fluctuation trajectories from a first attractor continued by the relaxation trajectories reaching the second attractor, for the pair (*Z*_+−_, *Z*_−+_). We omit the other pairs of attractors (*Z*_+−_, *Z*_−−_) and (*Z*_−−_, *Z*_−+_), because their optimal trajectories are simply straight lines.

Finally, we want to compute the optimal transition cost between any pair of basins (*Z*_*i*_, *Z*_*j*_). We observe that every relaxation trajectories starting on the common boundary of two basins stay on this boundary and converge to a unique saddle point inside: the assumption (A) of Theorem 5 is then verified. It follows from this theorem that the optimal trajectory between any basin *Z*_*i*_ and *Z*_*j*_ necessarily reaches *∂Z*_*i*_ ∩ *∂Z*_*j*_ on a saddle point, and then that the optimal transition cost is given by the trajectory which minimizes the cost among all those found previously between the attractor and the saddle points. We denote this optimal trajectory 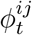. Its cost is explicitly described by the formula (29) (with *T* → ∞), which is then the optimal cost of transition between *Z*_*i*_ and *Z*_*j*_.

The LDP ensures that for all *δ, η* > 0, there exists *ε*′ such that for all *ε* ∈ (0, *ε*′), a random path 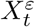 reaching *Z*_*j*_ from 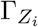 before 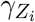, verifies: 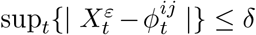 with probability larger than 1 − *η*. In other words, given a level of resolution *δ*, we could then theoretically find *ε* such that any trajectory of exit from *Z*_*i*_ to *Z*_*j*_ would be indistinguishable from trajectory 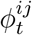 at this level of resolution. But in practice, the event 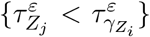 is too rare to be simulated directly for such *ε*.

We plot in Figure 7 two sets of random exit paths, simulated for two different *ε*, illustrating the fact that the probability of an exit path to be far from the optimal fluctuation trajectory decreases with *ε*.

**Figure 7:**
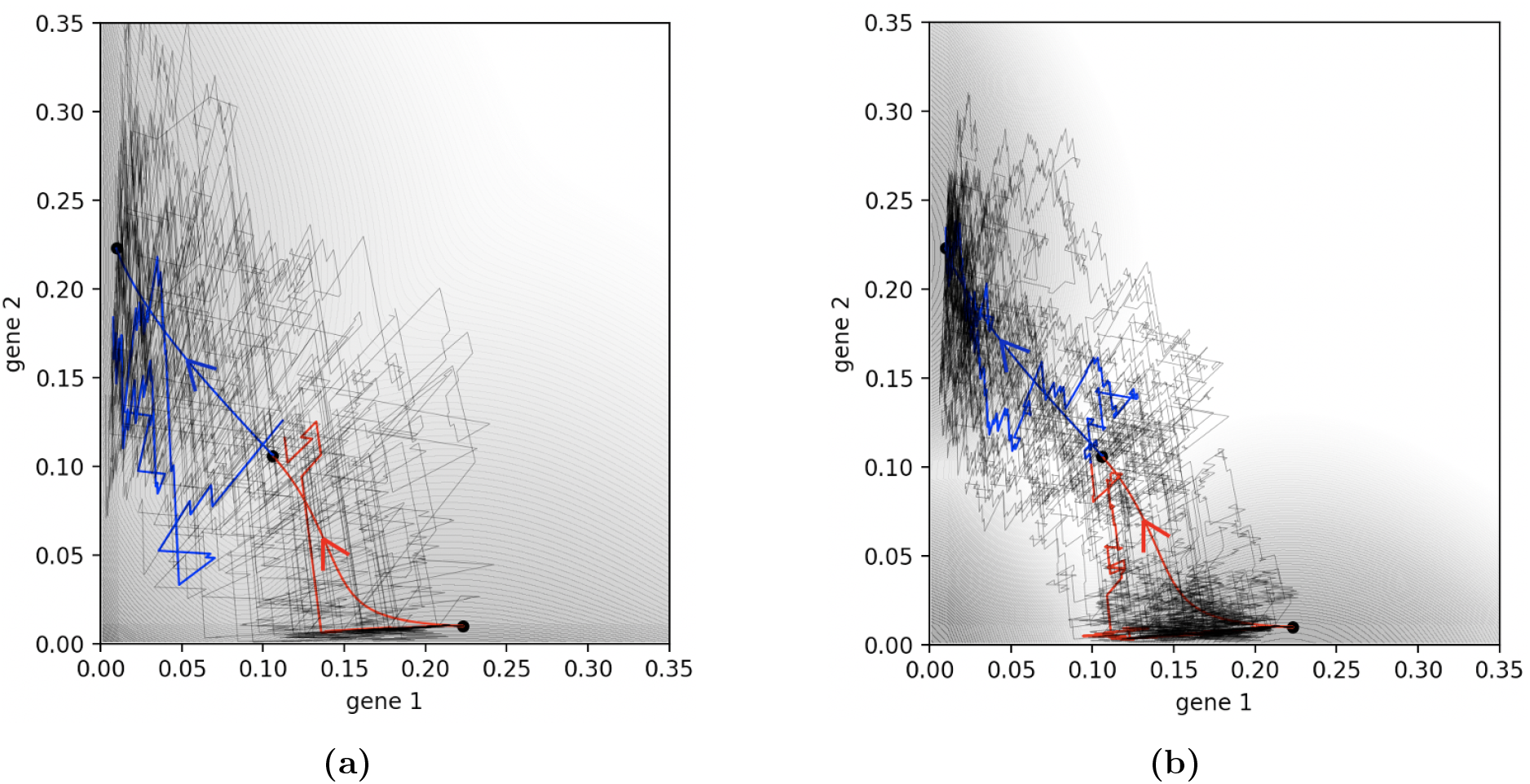
Comparison between the optimal trajectory of the figure 6 and 30 random paths conditioned on reaching, from a point of the boundary of the *R*-neighborhood of an attractor 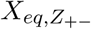, the *r*-neighborhood of a new attractor 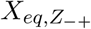 before the *r* neighborhood of the first attractor 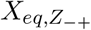, with *r* < *R*. We represent this comparison for 7a: *ε* = 1/7 and 7b: *ε* = 1/21. For each figure, one of these random paths is colored, separating the fluctuation and the relaxation parts.

### 5.2 Comparison between predictions and simulations

For each pair of basins (*Z*_*i*_, *Z*_*j*_), the expression (29) provides an approximation of the probability of the rare event 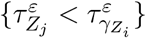, up to a prefactor, and the approximation (17) allows to deduce the associated transition rate. We plot in Figure 8 the evolution of these two estimations, as *ε* decreases, comparing respectively to the probabilities given by the AMS algorithm and the transition rates computed with a Monte-Carlo method. As in [50], we decide to plot these quantities in logarithmic scale. We observe that, knowing the prefactor, the Large deviations approximation is accurate even for *ε* > 0.1, and that induced transition rates are close to the ones observed with a Monte-Carlo method too. We represent in Figure 10b the variance of the estimator of the transition rates given by the AMS method.

**Figure 8:**
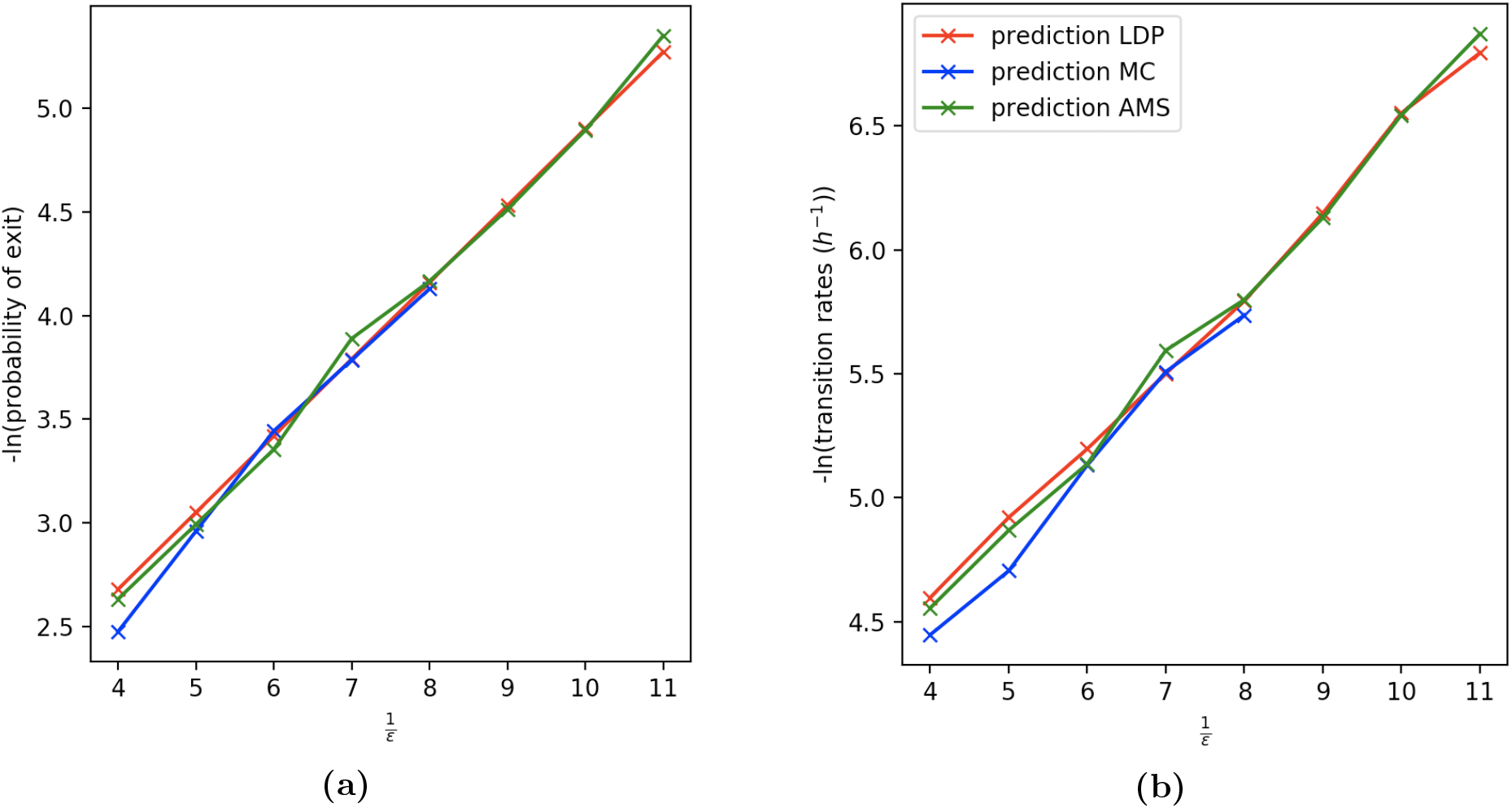
8a: Comparison between the probabilities (7) between the basins *Z*_+−_ and *Z*_−−_, in logarithmic scale, given by the Large deviations approximation (in red) and the AMS algorithm (in green). The prefactor is computed for *ε* = 1/8 and the red curve is then adjusted to fit the numerical results. The blue curve corresponds to the probabilities obtained with a Monte-Carlo method. 8b: Comparison between the transition rates between the basins *Z*_+−_ and *Z*_−−_, in logarithmic scale, given by the formula (8), where the probability (7) is given by the Large deviations approximation (in red) and the AMS algorithm (in green). The blue curve corresponds to the transition rates obtained with a Monte-Carlo method, by the formula (6). The quantities obtained by a Monte-Carlo method, in blue, are not represented after *ε* = 1/8 because the transition rates become too small to be efficiently computed.

We also remark that our analysis provides two ways of estimating the stationary measure of the discrete coarse-grained model. On the one hand, we can obtain a long-time proteins distribution of thousands of cells by simulating the PDMP system (1) from random initial conditions: by identifying each cell with a basin, as shown in Figure 9a, we can find a vector *μ*_*b*_ describing the ratio of cells belonging to each basin. When the number and length of the simulations are large enough, this vector *μ*_*b*_ should be a good approximation of the stationary measure on the basins. On the other hand, the transition rates allows to build the transition matrix *M* of the discrete Markov process on the basins, 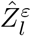, defined in Section 2.2. If the exponential approximation of the first passage time from every basin is accurate, then the stationary distribution on the basins should be well approximate by the unique probability vector such that *μ*_*z*_*M* = 0 (see Figure 9b).

**Figure 9:**
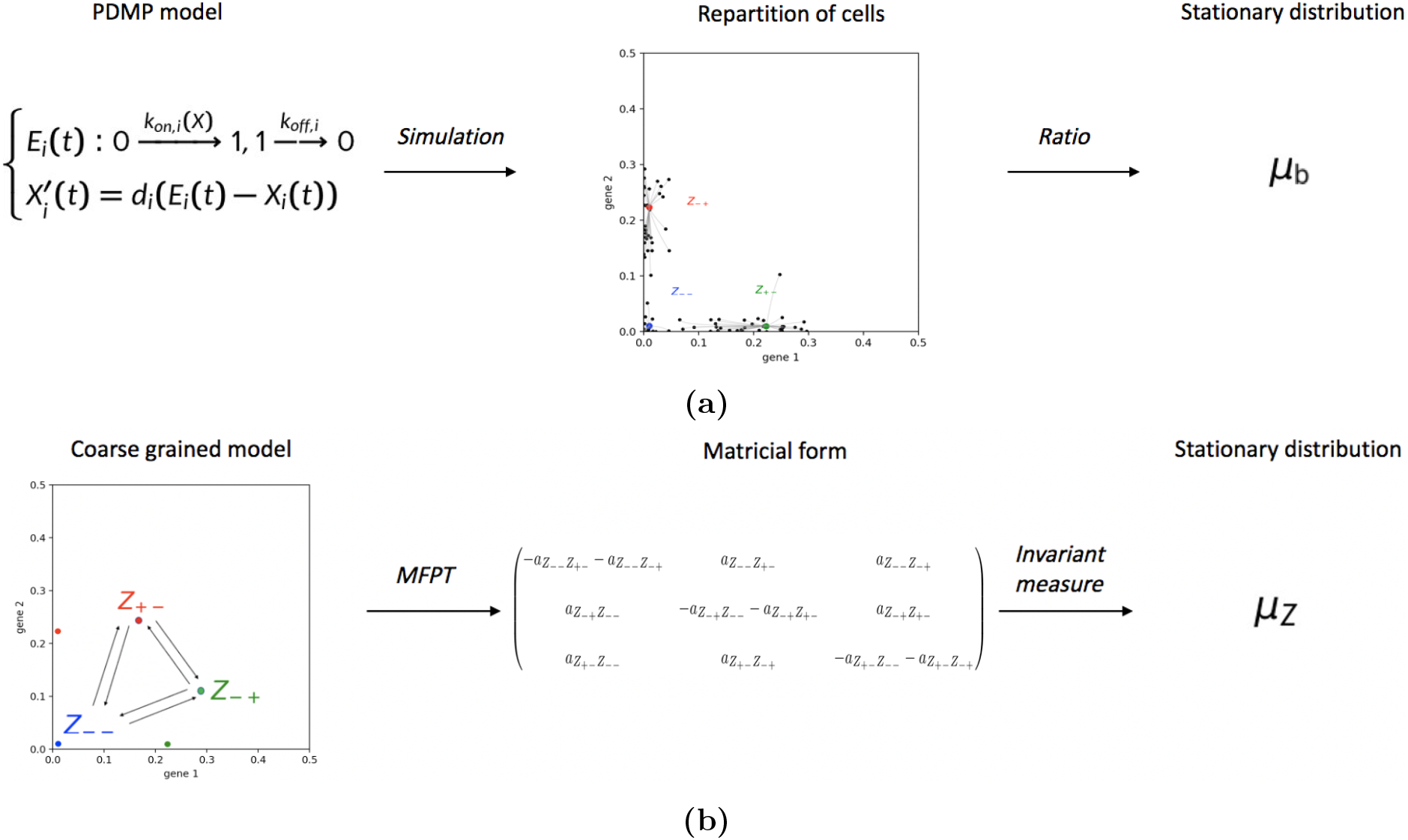
Comparison between the two methods for obtaining estimators of the stationary distributions on the basins: *μ*_*b*_ (9a) and *μ*_*z*_ (9b).

**Figure 10:**
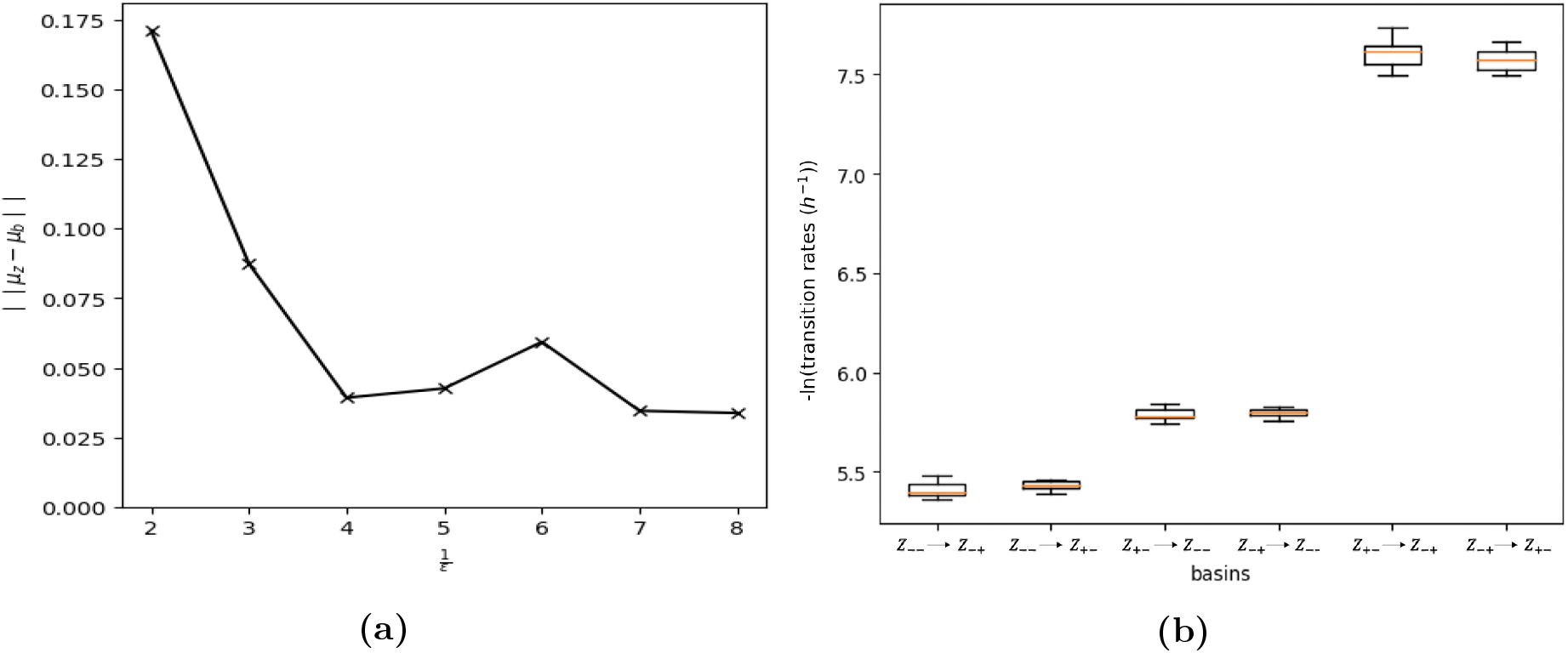
10a: The total variation of the difference between *μ*_*b*_ and *μ*_*z*_ as a function of *ε*^−1^. 10b: Boxplots representing the variation of the transition rates for 10 iterations of the method used in 10a, between each pair of basins for *ε* = 1/7.

Monte-Carlo methods for approximating the transition rates have a very high computational cost when *ε* is small. Thus, comparing these two stationary distributions appears as a good alternative for verifying the accuracy of the transition rates approximations. We plot in Figure 10a the evolution of the total variation distance between these two stationary distributions as *ε* decreases. We observe that the total variation is small even for realistic values of *ε*. The variance of the estimator *μ*_*b*_ is very small (given it is estimated after a time long enough) but the estimator *μ*_*z*_ accumulates all numerical errors coming from the estimators needed to compute the transition rates: this is likely to explain the unexpected small increases observed in this curve for *ε* = 1/6. We represent in Figure 10b the variance of the transition rates estimators between every pair of attractors used for estimating the distribution *μ*_*z*_ in Figure 10a, for *ε* = 1/7: as expected, this variance increases with the transition rates.

The similarity between the two distributions *μ*_*z*_ and *μ*_*b*_ seems to justify the Markovian approximation of the reduced process 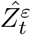 for small but realistic *ε*: at least for the toggle-switch network, the coarse-grained model, evolving on the basins of attractions seen as cellular types, describes accurately the complex behaviour of a cell in the gene expression space.

### 5.3 Applicability for more complex networks

It is in general very complex to find a solution *V* ∈ *C*1(Ω, ℝ) to the stationary Hamilton-Jacobi equation (25) which satisfies the condition (C) for general networks, when the number of genes is greater than 2. In order to apply the strategy developed in Section 4, for computing the cost of the optimal trajectories of transition between two basins, it would be then necessary to build a computational method for approximating such solution. Although the most common approach in this case consists in finding optimal trajectories without computing a potential (see [51] or [52] for more recent works), some methods have been recently built for SDEs model, like Langevin dynamics [53]. Such computational method for the PDMP system is beyond the scope of the article. However, we remark that even if there are no reasons for the trajectories satisfying the system (28) to be optimal when no function satisfying the relation (26) can be found, our computational method still allows to compute these trajectories, and we observe that they generally still bound the attractors and the saddle points of the deterministic system (4). Their costs can then be used as a proxy for the probabilities of the form (7): we observe in Figures 16b and 17b in Appendix J that for two non-symmetric networks of respectively 3 and 4 genes, our method still provides good results.

## 6 Discussion

Using the WKB approximation presented in Section 3.2.2 and the explicit formulas for the Hamiltonian and the Lagrangian detailed in Section 4.1, we are going now to analyze more precisely how the LDP for the proteins can be interpreted in regards to the dynamics of promoters, and we will see how two classical notions of energies can be interpreted in light of this analysis.

### 6.1 Correspondences between velocities and promoters frequency lead to energetic interpretations

The main idea behind the LDP principle for the PDMP system is that a slow dynamics on proteins coupled to the fast Markov chain on promoters rapidly samples the different states of *P*_*E*_ according to some probability measure 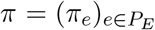. The value 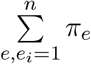 corresponds then to the parameter of the Bernoulli describing the random variable *E*_*i*_, and can be interpreted as the frequency of the promoter of gene *i*.

The point of view of [35] consisted in stating a LDP for the PDMP system by studying the deviations of *π* from the quasistationary distribution (3). The work of [29] consists in averaging the flux associated to the transport of each protein over the measure *π*, in order to build a new expression of this LDP which depends only on the protein dynamics. Its coupling with an Hamiltonian function through a Fenchel-Legendre transform allows to apply a wide variety of analytical tools to gain insight on the most probable behaviour of the process, conditioned on rare events. In this Section, we see how correspondences between these different points of view on the LDP shed light on the meaning of the Hamiltonian and Lagrangian functions and lead to some energetic interpretations.

#### 6.1.1 Correspondence between velocity and promoter frequency

Let us fix the time. The velocity field of the PDMP system, that we denote Φ, is a *n*-dimensional vector field, function of the random vectors *E, X*, which can be written for any *i* = 1, · · ·, *n*:

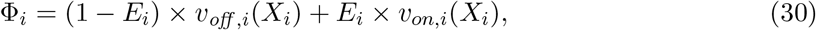

with the functions *v*_*off,i*_: *x*_*i*_ ↦ −*d*_*i*_*x*_*i*_ and *v*_*on,i*_: *x*_*i*_ ↦ *d*_*i*_(1 − *x*_*i*_) for any *x* ∈ Ω.

For all *i* = 1, · · ·, *n*, let 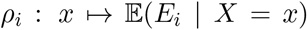 denote the conditional expectation of the promoter *E*_*i*_ knowing a protein vector *X*. As presented in Section 2.1, the quasistationary approximation identifies the vector field *ρ* to the invariant measure of the Markov chain on the promoter states.

For a given conditional expectation of promoters *ρ*, the vector field 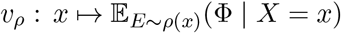 is defined for all *x* ∈ Ω by:

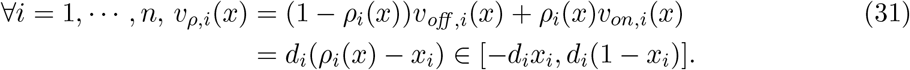

Denoting Ω^*v*^ the set of vector fields *v* continuous on Ω, such that for all *x* ∈ Ω, *v*(*x*) ∈ Ω^*v*^(*x*), we see that *v*_*ρ*_ ∈ Ω^*v*^. Conversely, the formula (31) can be inverted for associating to every velocity field *v* ∈ Ω^*v*^, characterizing the protein dynamics, a unique conditional expectation of promoters states knowing proteins, *ρ*_*v*_, which is the unique solution to the reverse problem 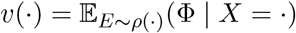, and which is defined by:

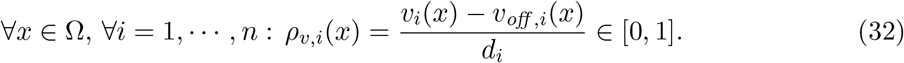

#### 6.1.2 Dynamics associated to a protein field

We detailed above the correspondence between any admissible velocity field *v* ∈ Ω^*v*^ and a unique vector field *ρ*_*v*_ describing a conditional expectation of promoters states knowing proteins. Moreover, the proof of Theorem 2 reveals that for any vector field *p*: Ω ↦ ℝ^*n*^, we can define a unique vector field *α*_*p*_: Ω ↦ (0, 1)^*n*^ by the expression (20).

As presented in Section 3.2.2, we denote *V* the leading order term of the Taylor expansion in *ε* of the function *S* defined in (13), such that the distribution of the PDMP system is defined at a fixed time *t*, and for all *e* ∈ *P*_*E*_, by 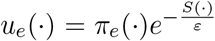, where *π*(*x*) is a probability vector in *S*_*E*_ for all *x* ∈ Ω.

On the one hand, we have seen in Section 3.2.2 that for all *x* ∈ Ω, the eigenvector *ζ*(*x,* ∇_*x*_*V* (*x*)) of the spectral problem (10) (for *p* = ∇_*x*_*V* (*x*)) corresponds to the leading order term of the Taylor expansion in *ε* of *π*(*x*). For all *i* = 1, · · ·, *n*, the quantity 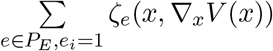 then represents the leading order approximation of the conditional expectation 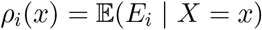. On the other hand, if we denote the gradient field *p* = ∇_*x*_*V* defined on Ω, we recall that for all 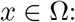 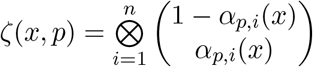. We then obtain:

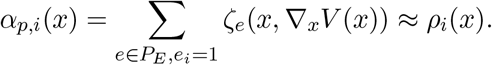

This interpretation of the vector *α*_*p*_, combined with the relation (32), allows us to state that the velocity field defined for all *x* ∈ Ω by 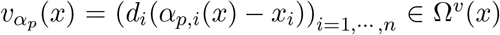 characterizes, in the weak noise limit, the protein dynamics associated to the proteins distribution 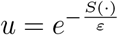. We see that the velocity field 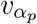 corresponds to the drift of the deterministic system (4) if and only if 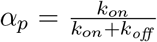, and then if and only if *p* = 0 (see Section 4.2). The gradient field *p* can be understood as a deformation of the deterministic drift, in the weak noise limit.

We recall that for all *p* ∈ ℝ^*n*^, we have from (21):

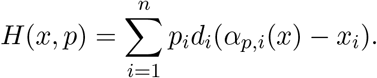

With the previous notations, the Lagrangian associated to a velocity field *v* can then be written on every *x* ∈ Ω as a function of *α*_*p*_ and *ρ*_*v*_:

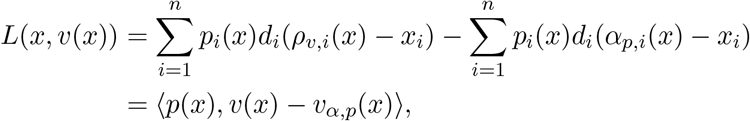

where *p*(*x*) = *p*_*v*_(*x*) is defined by the expression (24). Thus, we see that the duality between the Lagrangian and the Hamiltonian, that we intensively used in this article for analyzing the optimal trajectories of the PDMP system, and which is expressed through the relation (24) between the variables *v* and *p*, also corresponds to a duality between two promoters frequencies *ρ*_*v*_ and *α*_*p*_ associated to the velocity fields *v* and 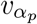.

The situation is then the following: for a given proteins distribution 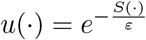 such that the first order approximation of *S* in *ε*, *V*, is differentiable on Ω, the velocity field *v* associated by duality to the gradient field *p* = ∇_*x*_*V*, and which characterizes a collection of optimal trajectories of the PDMP system (satisfying the system (16) associated to *V*) when *u* is the stationary distribution, does not correspond to the protein velocity 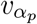 associated to the distribution *u* in the weak noise limit, except when the Lagrangian vanishes on (*x, v*). Alternatively speaking, the optimal trajectories associated to a distribution in the sense of Large deviations, characterized by the velocity field *v*, do not correspond to the trajectories expected in the weak noise limit, characterized by the velocity field 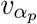. This is an important limit for developing a physical interpretation of the Hamiltonian system in analogy with Newtonian mechanics. However, the correspondence between promoters states distributions and velocity fields developed above leads us to draw a parallel with some notions of energy.

#### 6.1.3 Energetic interpretation

Following a classical interpretation in Hamiltonian system theory, we introduce a notion of energy associated to a velocity field:

##### Definition 4.

*Let us consider x* ∈ Ω *and v* ∈ Ω^*v*^. *The quantity E*_*v*_(*x*) = *H*(*x*, *p*_*v*_(*x*)) *is called the energy of the velocity field v on x, where p*_*v*_(*x*) *is defined by the expression* (24).

Interestingly, combining the expression of the Hamiltonian given in Theorem 2 with the expressions (24) and (32), the energy of a velocity *v* on every *x* ∈ Ω can be rewritten:

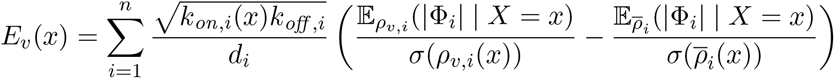

where for all *i* = 1, · · ·, *n*, Φ_*i*_ is the random variable defined by the expression (30), which follows, conditionally to proteins, a Bernoulli distribution of parameter *ρ*_*v,i*_, and 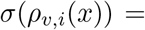 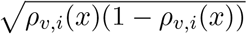 denotes its standard deviation.

Finally, we have 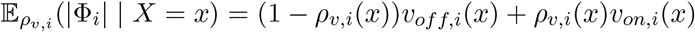, and 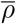 denotes the quasistationary distribution described in (3).

Formally, the energy of a promoter distribution can then be decomposed in two terms: a first term describing its velocity in absolute terms, scaled by its standard deviation, and a second term depending on the network. A high energy distribution on a point *x* is characterized by a fast and deterministic protein dynamics in regards respectively to the velocity of the quasistationary approximation on *x* and the standard deviation of its associated promoter distribution. We remark that this notion of energy does not depend on the proteins distribution, but only on the promoters frequency *ρ*_*v*_ around a certain location *x*. Depending on *x* only through the vector field *ρ*_*v*_ (and the functions *k*_*on,i*_), it is likely to be interpreted as the kinetic energy of a cell.

The potential 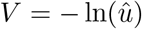, where 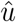 is the marginal on proteins of the stationary distribution of the stochastic process, is classically interpreted as a notion of potential energy, not depending on the effective promoter frequency. Apparently, this notion of energy is not related to the one described previously. Once again, the difficulty for linking these two notions of energy comes from the fact that the dynamics associated to the “momentum” *p* = ∇_*x*_*V*, which is characterized by the velocity field *v* defined by the formula (23), is not the same that the protein dynamics associated in the weak noise limit to the marginal distribution on proteins 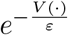, which is defined by the promoters frequency 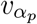.

### 6.2 Mixture model

The results of Section 5 lead us to consider the coarse-grained model as promising for capturing the dynamics of the metastable system, even for realistic *ε*. We are now going to introduce a mixture model which provides an heuristic link between the protein dynamics and the coarse-grained model, and appears then promising for combining both simplicity and ability to describe the main ingredients of cell differentiation process.

When *ε* is small, a cell within a basin *Z*_*j*_ ∈ *Z* is supposed to be most of the time close to its attractor: a rough approximation consists in identifying the activation rate of a promoter *e*_*i*_ in each basin by the dominant rate within the basin, corresponding to the value of *k*_*on,i*_ on the attractor. For any gene *i* = 1, · · ·, *n* and any basin *Z*_*j*_ ∈ *Z*, we can then consider:

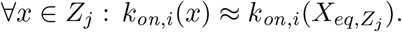

Combining this approximation of the functions *k*_*on,i*_ by their main mode within each basin with the description of metastability provided in Section 2.2, we build another process described by the 2*n +* 1-dimensional vector of variables (*Z*(*t*), *E*(*t*), *X*(*t*)), representing respectively the cell type, the promoter state and the protein concentration of all the genes (see Figure 11).

**Figure 11:**
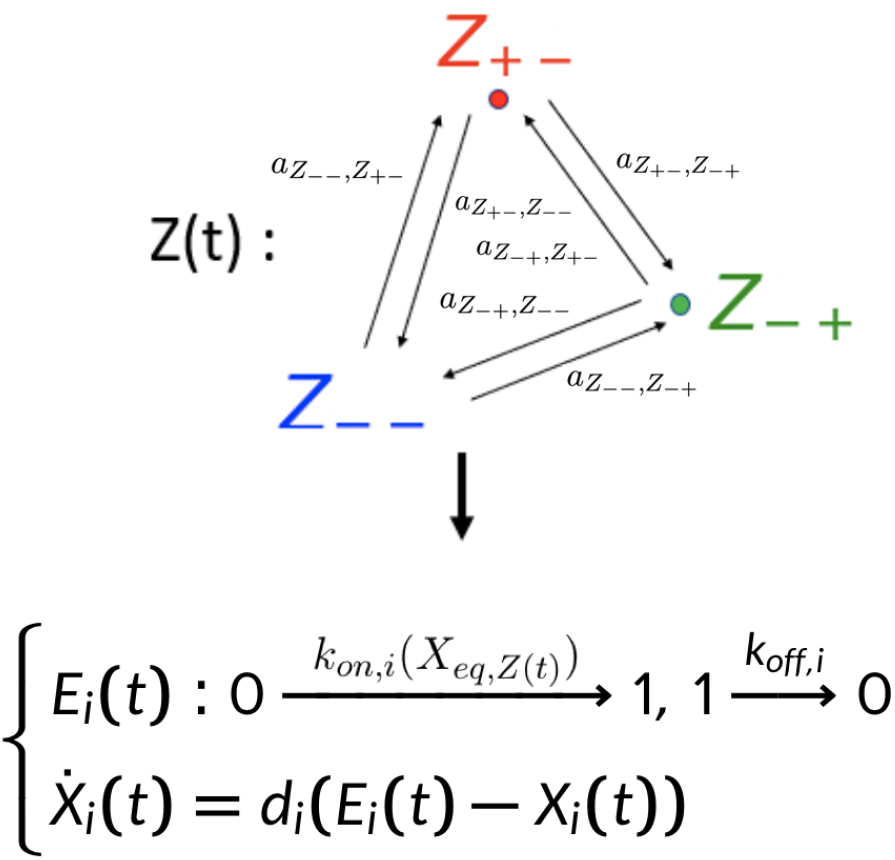
Weak noise approximate model. The Markov chain on the set of basins *Z* is here illustrated by the one corresponding to the toggle-switch network of Figure 3a.

Considering that the PDMP system spends in each basin a time long enough to equilibrate inside, we decide to approximate the distribution of the vector (*E*(*t*), *X*(*t*)) in a basin *Z*_*j*_ by its quasistationary distribution. It is then equivalent to the stationary distribution of a simple two states model with constant activation function, which is a product of Beta distributions [28]. Thus, the marginal on proteins of the stationary distribution of this new model, that we denote *u*, can be approximated by a mixture of Beta distributions:

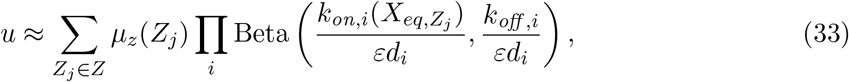

where *μ*_*z*_ is the stationary distribution of the Markov chain characterizing the coarse-grained model.

In that point of view, the marginal distribution on proteins of a single cell *X* is characterized by a hidden Markov model: in each basin *Z*_*j*_, which corresponds to the hidden variable, the vector *X* is randomly chosen under the quasistationary distribution 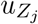 of the reduced process (*E, X* | *Z*_*j*_). This simplified model provides a useful analytical link between the proteins distribution of the PDMP system (depending on the whole GRN) and the coarse-grained model parameters.

This mixture also provides an approximation for the potential of the system on Ω:

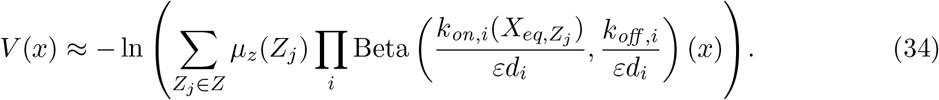

We remark that this new phenomenological model is a generalization of the local approximations of both the potential and the distribution within each basin that we have used for building the isocomittor surfaces and the score function of the AMS algorithm in Appendices F.3 and G.

### 6.3 One application for the mixture model

An interesting application for the mixture approximation presented in Section 6.2 is the computation of the potential energy of the system, as defined in the previous section. The potential energy of a population of cells *C* located on (*x*_*c*_)_*c*∈*C*_ can be approximated by the sum 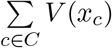, where *V* is defined by (34)

We represent in Figure 12 the evolution of the potential energy of a population of cells during the differentiation process, simulated from the PDMP system associated to the toggle-switch network presented in Appendix E. The population is initially centered on the attractor of the undifferentiated state *Z*_ _. We observe that the potential energy reaches a peak before decreasing.

**Figure 12:**
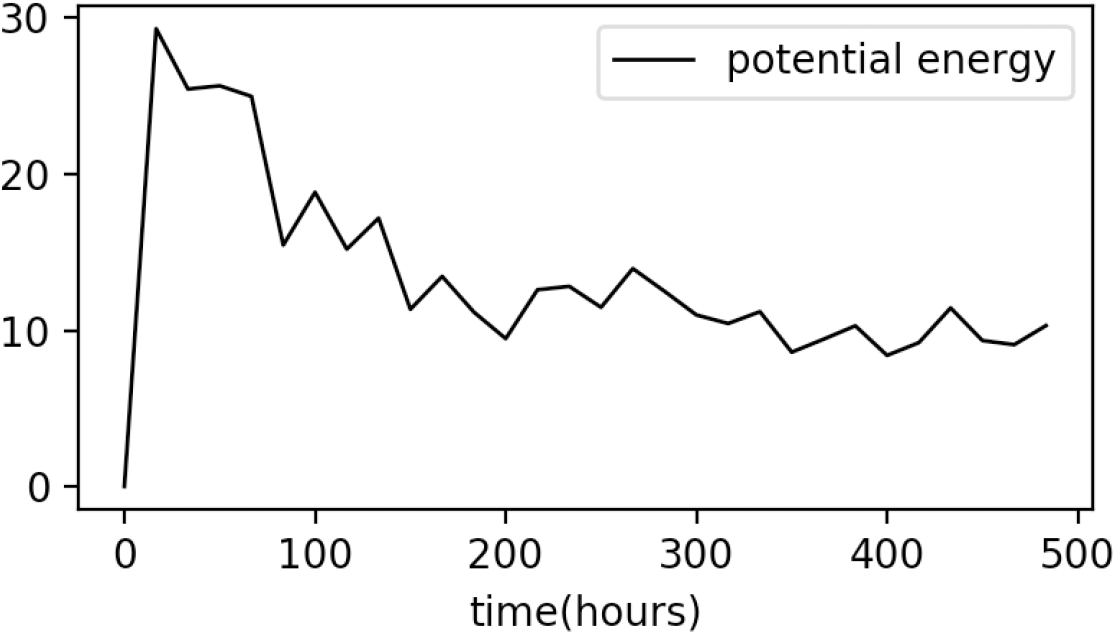
Evolution of the potential energy V of a population of 500 cells along the differentiation process.

We remark that in [54], the authors have revealed the universality of such feature during cell differentiation, for what they called the transcriptional uncertainty landscape, for many available single-cell gene expression data sets. This transcriptional uncertainty actually corresponds to the stationary potential *V* of our model, approximated for each cell from the exact stationary distribution of an uncoupled system of PDMPs (*i.e* with a diagonal interaction matrix). Although it cannot be formally linked to intracellular energetic spending yet, we can note that one of the authors recently described a peak in energy consumption during the erythroid differentiation sequence [55].

The mixture model also paves the way for interpreting non-stationary behaviours. Indeed, let us denote *μ*_*z,t*_ the distribution of the basins at any time *t*. The mixture distribution can be used as a proxy for non stationary distributions of a PDMP system:

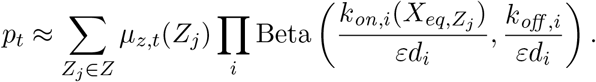

In that case, the only time-dependent parameters are the coordinates of the vector *μ*_*z,t*_ ∈ [0, 1]^*m*^ where *m* is the number of basins, and *μ*_*z,t*_ = *μ*_*z*_ if *t* is such that the stationary distribution is reached. The parameters 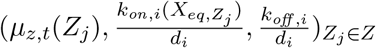 could be inferred from omics data at any time *t*, for example with an EM algorithm [56], [57].

## Conclusion

Reducing a model of gene expression to a discrete coarse-grained model is not a new challenge, ([26], [25]), and it is often hard to perform when the dimension is large. This reduction is closely linked to the notion of landscape through the quasipotential, the analysis of which has been often performed for non mechanistic models, where the random effects are considered as simple noise ([53], [22]), or for a number of genes limited to 2.

In this work, we propose a numerical method for approximating the transition rates of a multidimensional PDMP system modeling genes expression in a single cell. This method allows to compute these transition rates from the probabilities of some rare events, for which we have adapted an AMS algorithm. Although this method theoretically works for any GRN, the computation cost of the AMS algorithm may explode when both the number of genes increases and the scaling factor *ε* decreases.

In order to approximate these probabilities within the Large deviations context, we provided an explicit expression for the Hamiltonian and Lagrangian of a multidimensional PDMP system, we defined the Hamilton-Jacobi equation which characterizes the quasipotential, for any number of genes, and we provided the explicit expression of the associated variational problem which characterizes the landscape. We have deduced for some networks an analytical expression of the energetic costs of switching between the cell types, from which the transition rates can be computed. These approximations are accurate for a two-dimensional toggle-switch. We also verified that these analytical approximations seem accurate even for networks of 3 or 4 genes for which the energetic cost provided by the method is not proved to be optimal. However, testing the accuracy of this method for describing more complex networks would imply to build an approximate solution to the stationary Hamilton-Jacobi equation (25), which would be the subject of future works.

Finally, we have derived from the coarse-grained model a Beta-mixture model able to approximate the stationary behavior of a cell in the gene expression space. As far as we know, this is the first time that such an explicit link between a PDMP system describing cell differentiation and a non-Gaussian mixture model is proposed.

Altogether this work establishes a formal basis for the definition of a genetic/epigenetic landscape, given a GRN. It is tempting to now use the same formalism to assess the inverse problem of inferring the most likely GRN, given an (experimentally-determined) cell distribution in the gene expression space, a notoriously difficult task [58], [28].

Such random transitions between cell states have been recently proposed as the basis for facilitating the concomitant maintenance of transcriptional plasticity and stem cell robustness [59]. In this case, the authors have proposed a phenomenological view of the transition dynamics between states. Our work lays the foundation for formally connecting this cellular plasticity to the underlying GRN dynamics.

Finally our work provides the formal basis for the quantitative modelling of stochastic state transitions underlying the generation of diversity in cancer cells ([60], [61]), including the generation of cancer stem cells [62].

## Acknowledgments

This work was supported by funding from French agency ANR (SingleStatOmics; ANR-18-CE45-0023-03). We thank Ulysse Herbach for having highlighted the notions of main modes for the stochastic hybrid model of gene expression, and for critical reading of the manuscript. We would like to thank the referees and the associated editor for carefully reading our manuscript and for their constructive comments which helped improving the quality of the paper. We also thank all members of the SBDM and Dracula teams, and of the SingleStatOmics project, for enlightening discussions. We also thank the BioSyL Federation and the LabEx Ecofect (ANR-11-LABX-0048) of the University of Lyon for inspiring scientific events.

## Code availability

The code for reproducing the main figures of the article is available at https://gitbio.ens-lyon.fr/eventr01/jomb_reduction. It also contains the functions for the AMS algorithm, which is detailed in the appendix.

## A Mechanistic model and fast transcription reduction

We recall briefly the full PDMP model, which is described in details in [28], based on a hybrid version of the well-established two-state model of gene expression [30], [31] including both mRNA and protein production [63] and illustrated in Figure 13.

**Figure 13:**
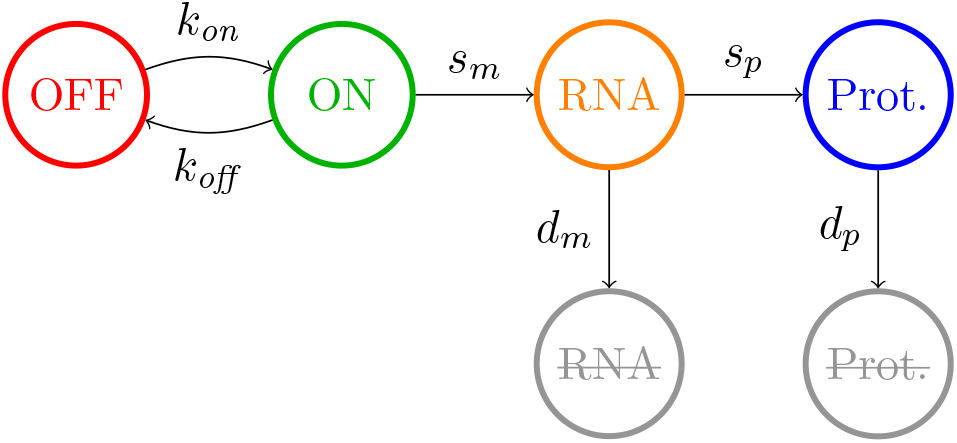
The two-states model of gene expression [28], [31].

A gene is described by the state of a promoter, which can be {*on, off*}. If the promoter is *on*, mRNAs will be transcripted with a rate *s*_*m*_ and degraded with a rate *d*_*m*_. If it is *off*, only mRNA degradation occurs. Translation of mRNAs into proteins happens regardless of the promoter state at a rate *s*_*p*_, and protein degradation at a rate *d*_*m*_. Neglecting the molecular noise of proteins and mRNAs, we obtain the hybrid model:

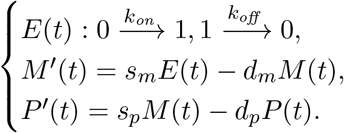

where (*E*(*t*), *M* (*t*), *P* (*t*)) denote respectively the promoter, mRNA and protein concentration at time *t*. As detailed in Section 1, the key idea is then to put this two-states model into a network by characterizing the jump rates of each gene by two specific functions *k*_*on,i*_ and *k*_*off,i*_, depending at any time on the protein vector *X*(*t*).

In order to obtain the PDMP system (1) that we use throughout this article, we exploit the two modifications that are performed in [28] to this mechanistic model. First, the parameters *s*_*m*_ and *s*_*p*_ can be removed to obtain a dimensionless model, from which physical trajectories can be retrieved with a simple rescaling.

Second, a scaling analysis leads to simplify the model. Indeed, degradation rates play a crucial role in the dynamics of the system. The ratio 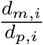 controls the buffering of promoter noise by mRNAs and, since *k*_*off,i*_ » *k*_*on,i*_, the ratio 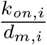 controls the buffering of mRNA noise by proteins. In line with several experiments [64] [65], we consider that mRNA bursts are fast in regard to protein dynamics, i.e 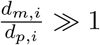 with 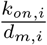 fixed. The correlation between mRNAs and proteins produced by the gene is then very small, and the model can be reduced by removing mRNA and making proteins directly depend on the promoters. We then obtain the PDMP system (1).

Denoting 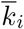 the mean value of the function *k*_*on,i*_, *i.e* its value where there is no interaction between gene *i* and the other genes, the value of a scaling factor 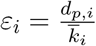 can then be decomposed in two factors: one describing the ratio between the degradation rates of mRNA and proteins, 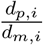, which is evaluated around 1/5 in [66], and one characterizing the ratio between promoter jumps frequency and the degradation rates of mRNA, 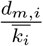. This last ratio is very difficult to estimate in practice. Assuming that it is smaller than 1, *i.e* that the mean exponential decay of mRNA when the promoter *E*_*i*_ is *off* is smaller than the mean activation rate, we can consider that *ε*_*i*_ is smaller than 1/5. Finally, for obtaining the model (1), we consider two typical timescales 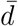 and 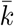, for the rates of proteins degradation and promoters activation respectively, such that for all genes *i*, 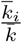 and 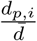 are of order 1 (when the disparity between genes is not too important). We then define 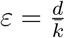.

## B Tensorial expression of the master equation of the PDMP system

We detail the tensorial expression of the master equation (2) for a two-dimensional network. We fix *ε* = 1 for the sake of simplicity.

The general form for the infinitesimal operator can be written:

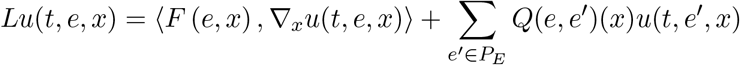

where *F* is the vectorial flow associated to the PDMP and *Q* the matrix associated to the jump operator.

A jump between two promoters states *e, e*′ is possible only if there is exactly one gene for which the promoter has a different state in *e* than in *e*′: in this case, we denote *e* ~ *e*′.

We have, for any *x*: *F* (*e, x*) = (*d*_0_(*e*_0_ − *x*_0_), · · ·, *d*_*n*_(*e*_n_ − *x*_*n*_))^*T*^. Then, for all *e* ∈ *P*_*E*_, the infinitesimal operator can be written:

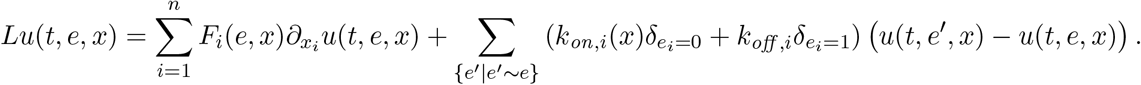

For a two-dimensional process (*n* = 2), there are four possible configurations for the promoter state: *e*_00_ = (0, 0), *e*_01_ = (0, 1), *e*_10_ = (1, 0), *e*_11_ = (1, 1). It is impossible to jump between the states *e*_00_ and *e*_11_. If we denote *u*(*t, x*) the four-dimensional vector: 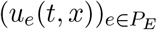, we can write the infinitesimal operator in a matrix form:

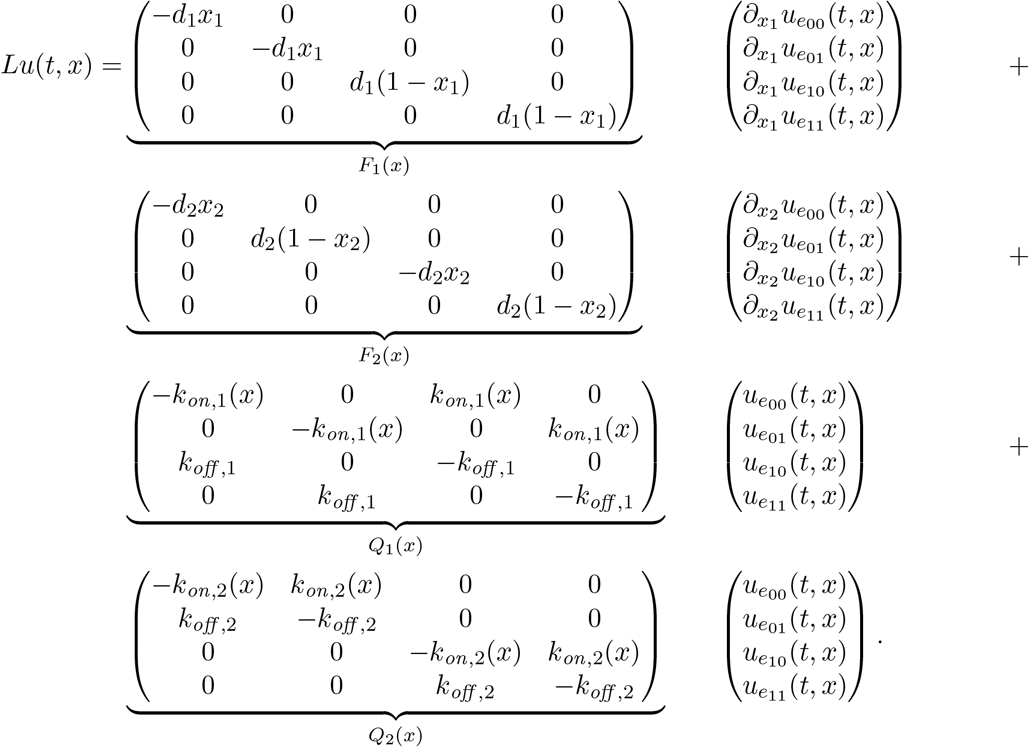

We remark that each of these matrices can be written as a tensorial product of the corresponding two-dimensional operator with the identity matrix:

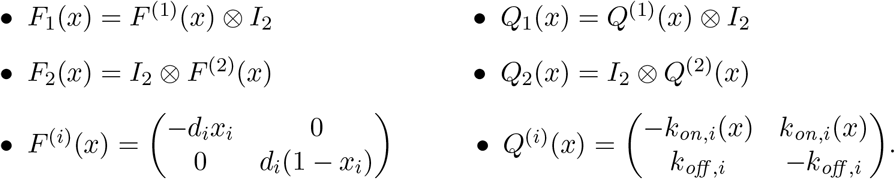

The master equation (2) is obtained by taking the adjoint operator of *L*:

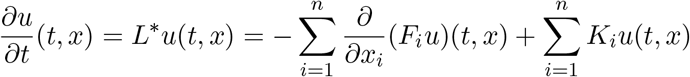

where *K*(*x*) = *Q*^*T*^ (*x*) is the transpose matrix of *Q*.

## C Diffusion approximation

## C.1 Definition of the SDE

In this section, we apply a key result of [36] to build the diffusion limit of the PDMP system (1). Let us denote 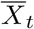 a trajectory satisfying the ODE system:

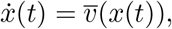

where 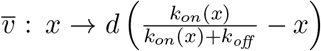 characterizes the deterministic system (4). We consider the process 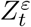 defined by:

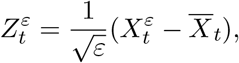

where *X*_*t*_*ε* verifies the PDMP system. Then, from the theorem 2.3 of [36] the sequence of processes 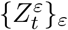 converges in law when *ε* → 0 to a diffusion process which verifies the system:

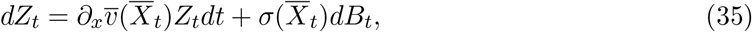

where *B*_*t*_ denotes the Brownian motion. The diffusion matrix Σ(*x*) = *σ*(*x*)*σ*^*T*^ (*x*) is defined by:

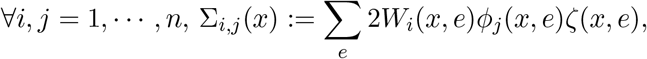

where ∀*e* ∈ *P*_*E*_, 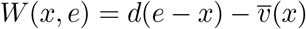, and *ϕ* is solution of a Poisson equation:

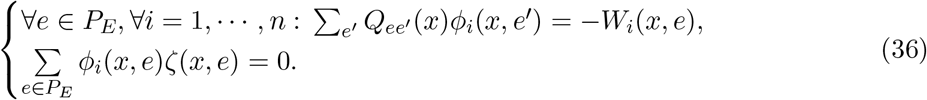

Let *ζ* be a probability vector in *S*_*E*_ representing the stationary measure of the jump process on promoters knowing proteins: ∀*x* ∈ Ω, *ζ*(*x,* ·)*Q*(*x*) = 0. We have: 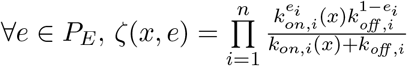.

It is straightforward to see that for all 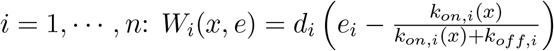. Then, let us define *ϕ* such that:

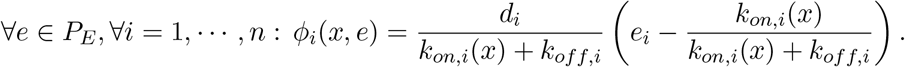

We verify that this vector *ϕ* is solution to the Poisson equation (36) for all *x*. The matrix Σ(*x*) is then a diagonal matrix defined by:

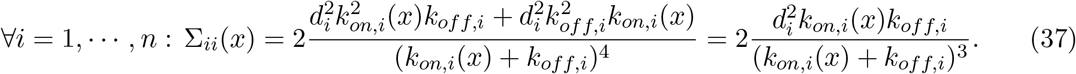

For all *x* ∈ Ω, the matrix *σ*(*x*) is then also diagonal and defined by:

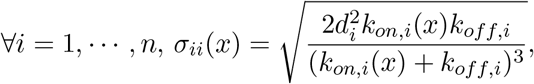

and we have defined all the terms of the diffusion limit (35).

## C.2 The Lagrangian of the diffusion approximation is a second-order approximation of the Lagrangian of the PDMP system

It is well known that the diffusion approximation satisfies a LDP of the form (12) [39]. The formula (37) allows to define the Lagrangian associated to this LDP, that we denote *L*_*d*_. From the theorem 2.1 of [39], we have:

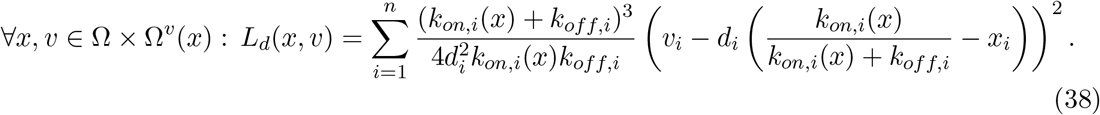

Note that for any fixed *x* ∈ Ω, *L*_*d*_(*x,* ·) is a quadratic function.

We recall that the Lagrangian associated to the LDP for the PDMP system, that we found in Theorem 3, is defined for all *x, v* ∈ Ω × Ω^*v*^(*x*) by:

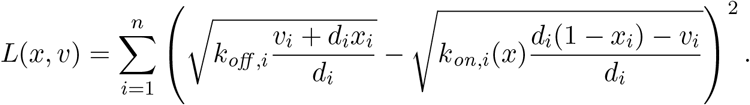

Expanding this Lagrangian with respect to *v* around 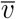 (the drift of the relaxation trajectories), we obtain:

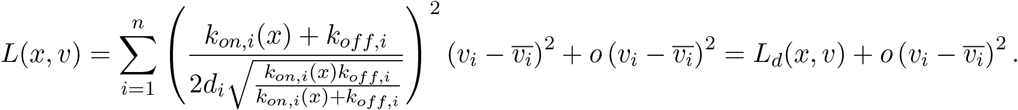

Thus, we proved that the Lagrangian of the diffusion approximation of the PDMP process corresponds to the two first order terms in 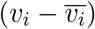 of the Taylor expansion of the real Lagrangian.

## D Example of interaction function

We recall that we assume that the vector *k*_*off*_ does not depend on the protein vector.

The specific interaction function chosen comes from a model of the molecular interactions at the promoter level, described in [28]:

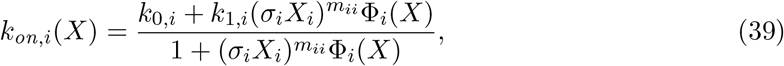

with:

- *k*_0,*i*_ the basal rate of expression of gene *i*,
- *k*_1,*i*_ the maximal rate of expression of gene *i*,
- *m*_*i,j*_ an interaction exponent, representing the power of the interaction between genes *i* and *j*,
- *σ*_*i*_ is the rescaling factor depending on the parameters of the full model including mRNAs,
- *θ* a matrix defining the interactions between genes, corresponding to a matrix with diagonal terms defining external stimuli, and
- 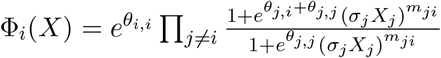

For a two symmetric two-dimensional network, we have for any *x* = (*x*_1_, *x*_2_) ∈ Ω:

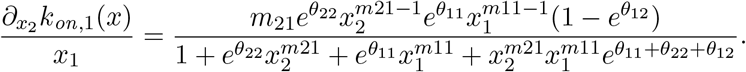

When *m*_11_ = *m*_22_ = *m*_12_ = *m*_21_ and *θ*_12_ = *θ*_21_, we have then for every *x* ∈ Ω:

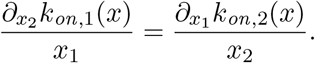

Thus, for all *x* ∈ Ω, when *d*_1_ = *d*_2_ we have:

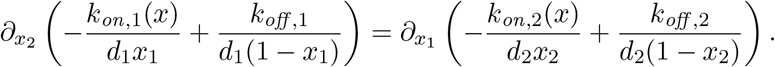

As a consequence, owing to the Poincaré lemma, there exists a function *V* ∈ *C*^1^(Ω, ℝ) such that the condition (26) is satisfied: one has

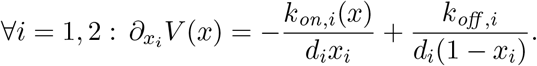

## E Description of the toggle-switch network

This table describes the parameters of the symmetric two-dimensional toggle-switch used all along the article. These values correspond to the parameters used for the simulations. The rescaling in time by the parameter scale 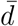, for the model presented in Section 1, corresponds to divide every *k*_0,*i*_, *k*_1,*i*_, *d*_*i*_ by 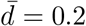. The mean values 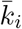 and *d*_*i*_ are then, as expected, of order 1 for every gene *i*.

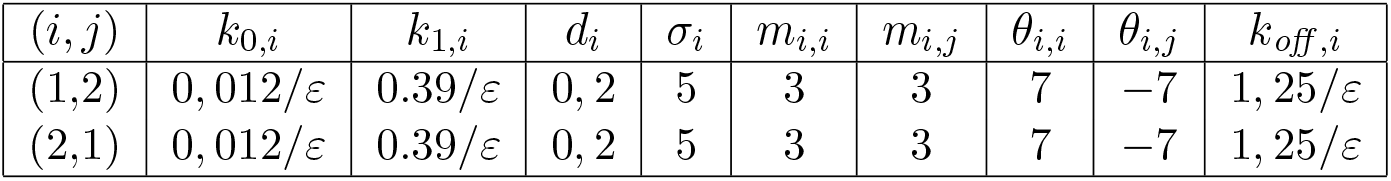

## F Details on the approximation of the transition rate as a function of probability (7)

In this section, we adapt the method developed in [40] to justify the formula (8) provided in Section 3.1, which approximate for every pair of basins (*Z*_*i*_, *Z*_*j*_) the transition rate *a*_*ij*_ as a function of the probability (7).

## F.1 General setting

Let us consider *r, R* such that 0 < *r* < *R*, we recall that 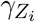 and 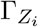 denote respectively the *r*-neighborhood and the *R*-neighborhood of the attractor 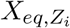. Let us consider a random path 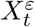 of the PDMP system, with initial condition 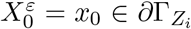. We define the series of stopping times 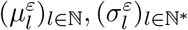 such that 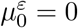 and for all *l* ∈ ℕ*:

- 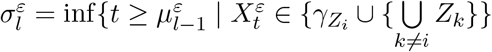
- 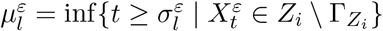

We then define 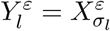. If 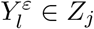, we set 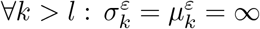 and the chain 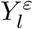 stops.

**Figure 14:**
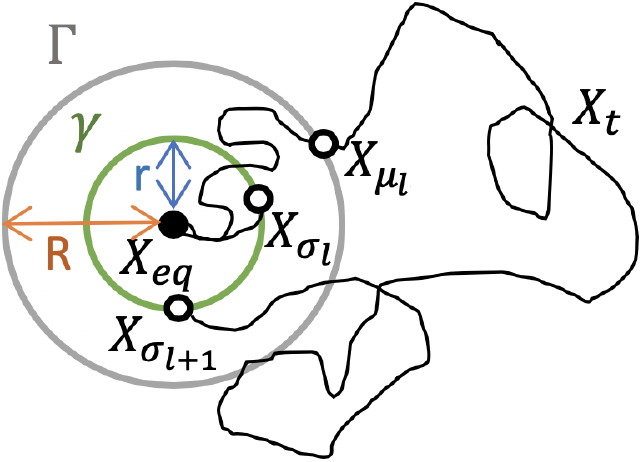
Illustration of the stopping times *σ*_*l*_ and *μ*_*l*_ describing respectively the *l*^*th*^ entrance of a random path *X*_*t*_ in a *r*-neighborhood *γ* of an attractor *X*_*eq*_, and its *l*^*th*^ exit from a *R*-neighborhood Γ.

From the formula (6) characterizing the transition rates, we can write:

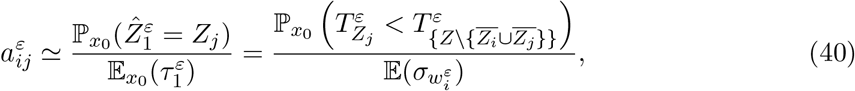

where we define the random variable: 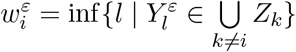.

Let us denote 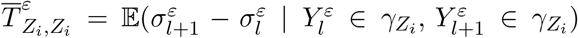. We can make the following approximation:

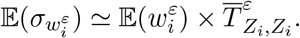

Indeed, the quantity on the left hand side is close to the mean number of attempts for reaching, from 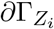, a basin *Z*_*k*_, *k* ≠ *i*, before 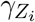, which is equal to 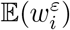, multiplied by the mean time of each attempt (knowing that at each step *l*, 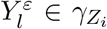), which is exactly 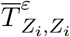. We should add the mean time for reaching *∂Z*_*j*_ from 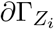 at the last step, but it is negligible when the number of attempts is large, which is the case in the small noise limit.

## F.2 Method when 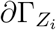 is reduced to a single point

We consider the case when 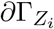 is reduced to a single point. It can happen for example when we consider only one gene (Ω = (0, 1)) and when the attractor 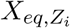 is located at a distance smaller than *r* from one of the boundaries of the gene expression space 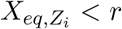 or 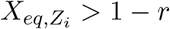. In such situation, a random path crosses necessarily the same point *x*_0_ to both exit 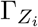 and come back to 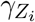 (if it does not reach a basin *Z*_*j*_ before): the Markov property of the PDMP process then justifies that the quantities 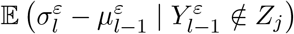 and 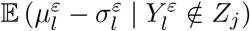 do not depend of *l*. Then 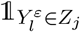 behaves like a discrete homogeneous Markov chain with two states, 1 being absorbing.

Let us define a second random variable 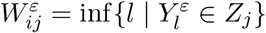. The homogeneity of the Markov chain 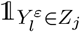 ensures that 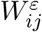 follows a geometric distribution. Its expected value is then the reverse of the parameter of the geometric law, *i.e*: 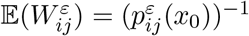.

Moreover, it is straightforward to see that from the same reasoning applied to any *Z*_*k*_, *k* ≠ *i*:

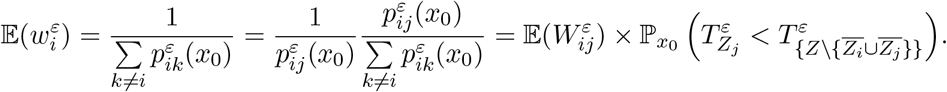

Thus, from (40) we can approximate the transition rate by the formula (8).

## F.3 Method in the general case

The difficulty for generalizing the approach described above, when 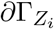 is not reduced to a single point, is to keep the Markov property, which has been used to cut the trajectories into pieces. Heuristically, the same argument which led us to approximate the PDMP system by a Markov jump process can be used to justify the asymptotic independence on *l* of the quantity 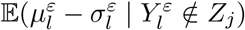: for *ε* ≪ 1, any trajectory starting on 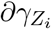 will rapidly loose the memory of its starting point after a mixing time within 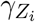. But it is more complicated to conclude on the independence from *l* of the quantity 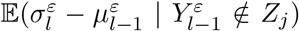, which may depend on the position of 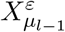 when the gene expression space is multidimensional.

We introduce two hypersurfaces 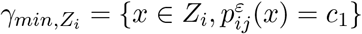 and 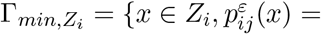 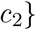, where *c*_1_ < *c*_2_ are two small constants. We substitute to the squared euclidean distance, used for characterizing the neighborhood 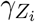 and 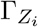, a new function based on the probability of reaching the (unknown) boundary: ∀*x, y* ∈ *Z*_*i*_, 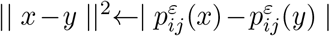. The function 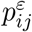 is generally called committor, and the hypersurfaces 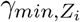 and 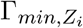 isocommittor surfaces. The committor function is not known in general; if it was, employing a Monte-Carlo method would not be necessary for obtaining the probabilities (7). However, it can be approximated from the potential of the PDMP system within each basin, defined in the equilibrium case by the well-known Boltzman law: 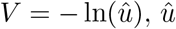 being the marginal on proteins of the stationary distribution of the process. Indeed, for reasons that are precisely the subject of Section 3.2 (studied within the context of Large deviations), the probability 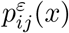 is generally linked in the weak noise limit to the function *V* by the relation:

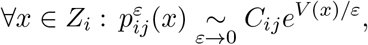

where *C*_*ij*_ is a constant specific to each pair of basins (*Z*_*i*_, *Z*_*j*_). We remark that when *ε* is small, a cell within a basin *Z*_*j*_ ∈ *Z* is supposed to be most of the time close to its attractor: a rough approximation could lead to identify the activation rate of a promoter *e*_*i*_ in each basin by the dominant rate within the basin, corresponding to the value of *k*_*on,i*_ on the attractor. For any gene *i* = 1, · · ·, *n* and any basin *Z*_*j*_ ∈ *Z*, we can then approximate:

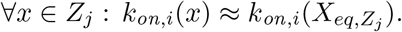

Under this assumption, the stationary distribution of the process is close to the stationary distribution of a simple two states model with constant activation function, which is a product of Beta distributions [28]. We then obtain an approximation of the marginal on proteins of the stationary distribution within each basin *Z*_*j*_:

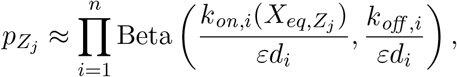

By construction, this approximation is going to be better in a small neighborhood of the attractor 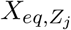. Thus, this expression provides an approximation of the potential *V* around each attractor 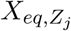:

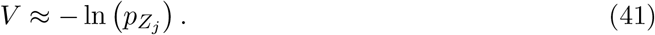

In every basin *Z*_*i*_, and for all *x* ∈ *Z*_*i*_ close to the attractor, the hypersurfaces where 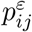 is constant will be then well approximated by the hypersurfaces where the explicitly known function 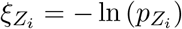 is constant.

For each attractor 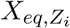, we can then approximate the two isocommittor surfaces described previously:

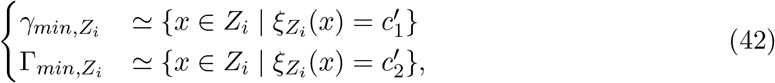

where *c*′_1_ and *c*′_2_ are two constants such that 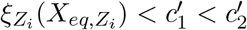.

We then replace 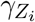 and 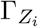 by, respectively, 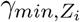 and 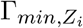 in the definitions of the stopping times 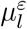 and 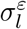 provided in Section F.1. From the proposition 1. of [40], we obtain that, as in the simple case described in Section F.2, 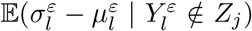 is independent of *l*. Defining 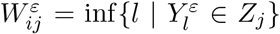, the definition of 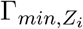 allows to ensure that 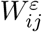 does not depend on the point *x*_0_ of 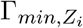 which is crossed at each step 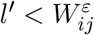. This random variable follows then a geometric distribution, with expected value 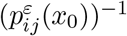, and we can derive an expression of the form (8).

## G AMS algorithm

We use an Adaptive Multilevel Splitting algorithm (AMS) described in [41]. The algorithm provides for every Borel sets (*A, B*) an unbiased estimator of the probability:

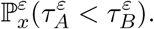

It is supposed that the random process attains easily *A* from *x*, more often than *B*, called the target set.

The crucial ingredient we need to introduce is a score function *ξ*(·) to quantify the adaptive levels describing how close we are from the target set *B* from any point *x*. The variance of the algorithm strongly depends on the choice of this function.

The optimal score function is the function 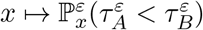 itself, called the committor which is unknown. It is proved, at least for multilevel splitting algorithms applied to stochastic differential equations in [67], [68], that if a certain scalar multiplied by the score function is solution of the associated stationary Hamilton-Jacobi equation, where the Hamiltonian comes from the Large deviations setting, the number of iterations by the algorithm to estimate the probability in a fixed interval confidence grows sub-exponentially in *ε*.

For the problem studied in this article, for every basin *Z*_*j*_ ∈ *Z*, we want to estimates probabilities substituting *A* to *γ*_*j*_ and *B* to another basin *Z*_*k*_, *k* ≠ *j*. Using the approximation of *V* given by the expression (34), we obtain the following score function, up to a specific constant specific to each basin:

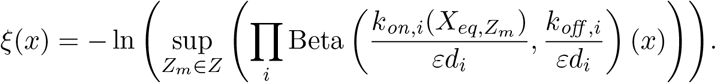

We remark that this last approximation allows to retrieve the definition of the local potential (41) defined on Appendix F.3, when the boundary of the basins are approximated by the leading term in the Beta mixture. The approximation is justified by the fact that for small *ε*, the Beta distributions are very concentrated around their centers, meaning that for every basin *Z*_*k*_ ∈ *Z, k* ≠ *j*:

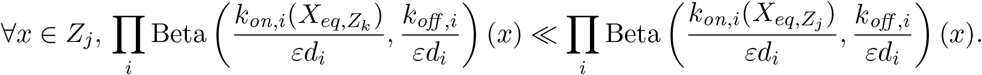

We supposed that ∀*Z*_*j*_ ∈ *Z, μ*_*z*_(*Z*_*j*_) > 0, where *μ*_*z*_ denotes the distributions on the basins. This is a consequence of the more general assumption that the stationary distribution of the PDMP system is positive on the whole gene expression space, which is necessary for rigorously deriving an analogy of the Gibbs distribution for the PDMP system (see Section 3.2.2).

We modify the score function to be adapted for the study of the transitions from each basin *Z*_*j*_ to *Z*_*k*_, *k* ≠ *j*:

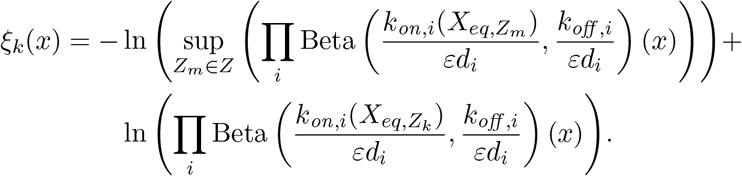

This function is specific to each transition to a basin *Z*_*k*_ but defined in the whole gene expression space. We verify: *ξ*_*k*_(*x*) ≤ 0 if *x* ∈ Ω \ *Z*_*k*_ and *ξ*_*k*_(*x*) = 0 if *x* ∈ *Z*_*k*_. We use *ξ*_*k*_ as the score function for the AMS algorithm.

In order to estimate probabilities of the type 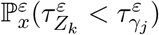 for *x* ∈ *Z*_*j*_, we need to approximate the boundaries of the basins of attraction, which are unknown. For this sake, we use again the approximate potential function *ξ* ≈ *V* to approximate the basins only from the knowledge of their attractor:

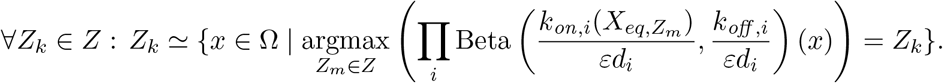

We use the Adaptative Multilevel Splitting algorithm described in Section 4 of [41], with two slight modifications in order to take into account the differences due to the underlying model and objectives:

- First, a random path associated to the PDMP system does not depend only on the protein state but is characterized at each time *t* by the 2n-dimensional vector: (*X*_*t*_, *E*_*t*_). For any simulated random path, we then need to associate an initial promoter state. However, we know that in the weak noise limit, for a protein state close to the attractor of a basin, the promoter states are rapidly going to be sampled by the quasistationary distribution: heuristically, this initial promoter state will not affect the algorithm. We decide to initially choose it randomly under the quasistationary distribution. For every *x*_0_ ∈ Γ_*min,j*_ beginning a random path in a basin *Z*_*j*_, we choose for the promoter state of any gene *i*, 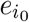, following a Bernoulli distribution:

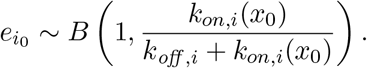
- Compared with [41], an advanced algorithm is used to improve the sampling of the entrance time in a set *γ*_*min,j*_. In practice timestepping is required to approximate the protein dynamics, and it may happen that the exact solution enters *γ*_*min,j*_ between two time steps, whereas the discrete-time approximation remains outside *γ*_*min,j*_. We propose a variant of the algorithm studied in [69] for diffusion processes, where a Brownian Bridge approximation gives a more accurate way to test entrance in the set *γ*_*min,j*_.

In the case of the PDMP system, we replace the Brownian Bridge approximation, by the solution of the ODE describing the protein dynamics: considering that the promoter state *e* remains constant between two timepoints, the protein concentration of every gene *i*, *x*_*i*_(*t*) is a solution of the ODE: 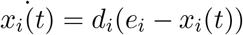, which implies:

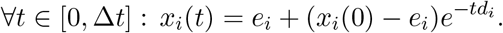

We show that for one gene, the problem can be easily solved. Indeed, let us denote 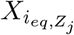 the *i*^*th*^ component of the vector 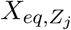. The function: 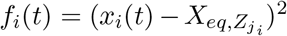 is differentiable and its derivative

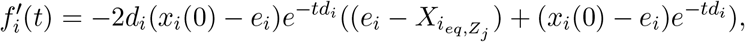

vanishes if and only if 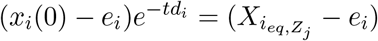, *i.e* when

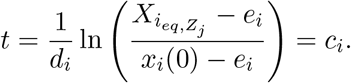

Then, if *c*_*i*_ ≤ 0 or *c*_*i*_ ≥ Δ*t*, the minimum of the squared euclidean distance of the *i*-th coordinate of the path to the attractor is reached at one of the points *x*_*i*_(0) or *x*_*i*_(Δ*t*). If 0 ≤ *c*_*i*_ ≤ Δ*t*, the extremum is reached at *x*_*i*_(*c*_*i*_). This value, if it is a minimum, allows us to determine if the process has reached any neighborhood of an attractor 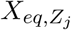 between two timepoints.

For more than one gene, the minimum of the sum: 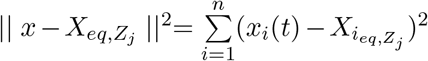 is more complicated to find. If for all *i* = 1, · · ·, *n*, *d*_*i*_ = *d*, which is the case of the two-dimensional toggle-switch studied in Section 5, the extremum can be explicitly computed:

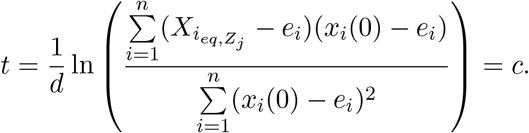

But we recall that for more than one gene, the set of interest is the isocommittor surface *γ*_*min,j*_ and not a neighborhood *γ*_*j*_. An approximation consists in identifying *γ*_*min,j*_ to the *r*-neighborhood of 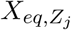, where *r* is the mean value of 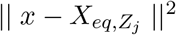 for *x* ∈ *γ*_*min,j*_.

If the parameters *d*_*i*_ are not all similar, we have to make the hypothesis that the minimum is close to the minimum for each gene. In this case, we just verify that for any gene *i*, the value of the minimum *x*_*i*_(*c*_*i*_) for every gene is not in the set 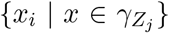: if it is the case for one gene, we consider that the process has reached the neighborhood 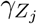 of the basins *Z*_*j*_ between the two timepoints.

## H Proofs of Theorems 4 and 5

First, we recall the theorem of characteristics applied to Hamilton-Jacobi equation [70], which states that for every solution *V* ∈ *C*^1^(Ω, ℝ) of (25), the system (16) associated to *V*

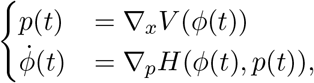

is equivalent to the following system of ODEs on (*x, p*) ∈ Ω × ℝ^*n*^, for *x*(0) = *ϕ*(0) and *p*(0) = Δ_*x*_*V* (*x*(0)):

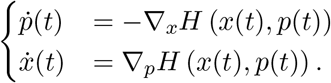

A direct consequence of this equivalence with an ODE system is that two optimal trajectories associated to two solutions of the stationary Hamilton-Jacobi equation cannot cross each other with the same velocity. We then have the following lemma:

#### Lemma 2.

*Let V*_1_ *and V*_2_ *be two solutions of* (25) *in C*^1^(Ω, ℝ).

*For any trajectories* 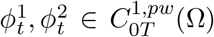 *solutions of the system* (16) *associated respectively to V*_1_ *and V*_2_*, if there exists t* ∈ [0, *T*] *such that ϕ*^1^(*t*) = *ϕ*^2^(*t*) *and* 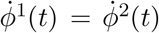, *then one has ϕ*^1^(*t*) = *ϕ*^2^(*t*) *for all t* ∈ [0, *T*].

This corollary is important for the two first items of the proof of Theorem 4:

#### Corollary 1.

*For any solution V* ∈ *C*^1^(Ω, ℝ) *of* (25) *and any trajectory* 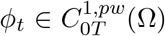 *satisfying the system* (16) *associated to V, we have the equivalence:*

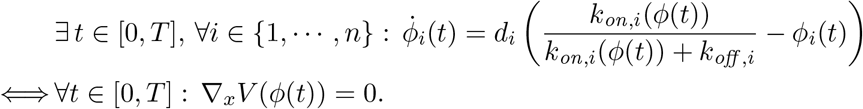

*Proof.* We recall that the relaxation trajectories correspond to trajectories satisfying the system (16) associated to a constant function *V*, *i.e* such that ∇_*x*_*V* = 0 on the whole trajectory. At any time *t*, the correspondence between any velocity field *v* of Ω^*v*^ and a unique vector field *p*, proved in Theorem 3 with the relation (24), allows to ensure that:

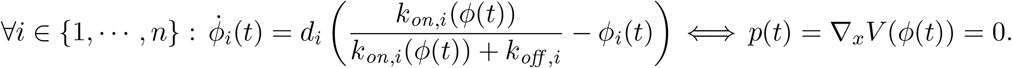

The lemma 2 ensures that any trajectory which verifies the same velocity field than a relaxation trajectory at a given time *t* is a relaxation trajectory: we can then conclude.

Finally, the following lemma is important for the first item of the proof of Theorem 4:

#### Lemma 3.

∀*i* ∈ {1, · · ·, *n*}, ∀*x* ∈ Ω *we have:*

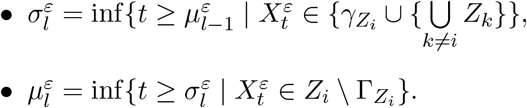

*Proof.* We have seen in the proof of Theorem 2 that for all *i* = 1, · · ·, *n* and for all *x* ∈ Ω, *H*_*i*_(*x,* ·) is strictly convex, and that *H*_*i*_(*x, p*_*i*_) → ∞ as *P*_*i*_ → ±∞. Moreover, *H*_*i*_(*x, p*_*i*_) vanishes on two points 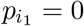 and 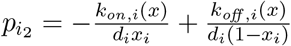 inside ℝ.

Then, the min on *p*_*i*_ is reached on the unique critical point 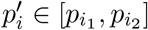, and we have: 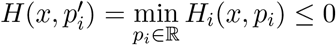

Finally:

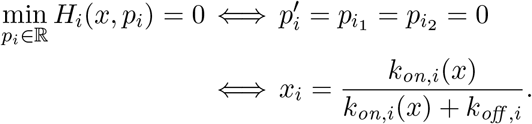

We now prove the theorem 4.

*Proof of Theorem 4.(i).* We consider a trajectory *ϕ*_*t*_ satisfying the system (16) associated to *V*. We recall that the Fenchel-Legendre expression of the Lagrangian allows to state that the vector field *p* associated to the velocity field 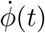 by the relation (24) is precisely *p* = ∇_*x*_*V* (*ϕ*(*t*)). When *V* is such that *H*(·, ∇_*x*_*V* (·)) = 0 on Ω, we have then for any time *t*:

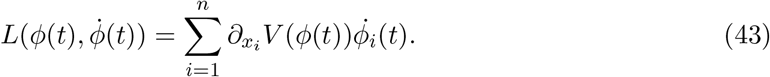

We recall that from Theorem 3:

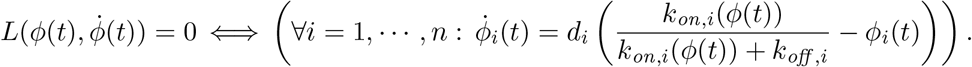

From this and (43), we deduce that for such an optimal trajectory:

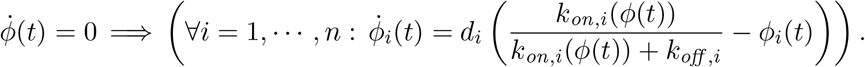

The velocity field then vanishes only at the equilibrium points of the deterministic system. Conversely, we recall that for such trajectory we have for any *t*:

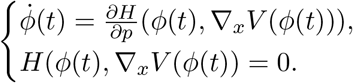

Assume that for all 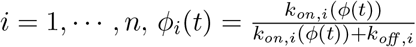. Then, by Lemma 3, we have: 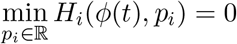 for all *i*.

Thereby, *H*(*ϕ*(*t*), ∇ _*x*_*V* (*ϕ*(*t*))) = 0 if and only if for all *i H*_*i*_(*ϕ*(*t*), 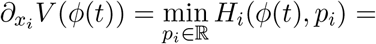 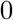, which implies: 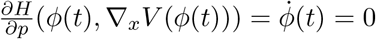. The lemma is proved.

*Proof of Theorem 4.(ii).* From the Corollary 1, if ∇_*x*_*V* (*ϕ*(*t*)) = 0, the trajectory is a relaxation trajectory, along which the gradient is uniformly equal to zero. The condition (C) implies that it is reduced to a single point: 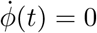. Conversely, with the same reasoning that for the proof of (i):

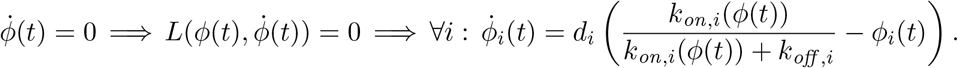

We recognize the equation of a relaxation trajectory, which implies: ∇_*x*_*V* (*ϕ*(*t*)) = 0. Thus, for any optimal trajectory satisfying the system (16) associated to a solution *V* of the equation (25) satisfying the condition (C), we have for any time *t*:

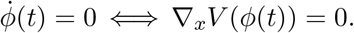

From the equivalence proved in (i), the condition (C) then implies that the gradient of *V* vanishes only on the stationary points of the system (4).

*Proof of Theorem 4.(iii).* As a consequence of (ii), for any optimal trajectory *ϕ*_*t*_ associated to a solution *V* of (25) which satisfies the condition (C), we have for all *t* > 0:

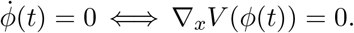

Then, if there exists *t* > 0 such that 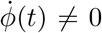, it cannot be equal to the drift of a relaxation trajectory, defined by the deterministic system (4), which is known to be the unique velocity field for which the Lagrangian vanishes (from Theorem (3)). Then it implies:

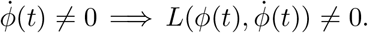

The relation (43), combined to the fact that the Lagrangian is always nonnegative allow to conclude:

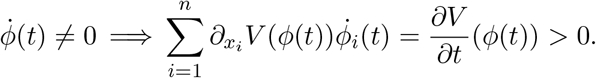

Thus, the function *V* strictly increases on these trajectories.

Furthermore, on any relaxation trajectory *ϕ*_*r*_(*t*), from the inequality (15) we have for any times *T* 1 < *T* 2:

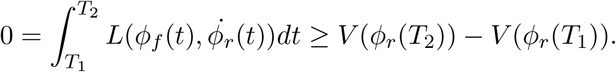

The equality holds between *T* 1 and *T* 2 if and only if for any *t* ∈ [*T* 1, *T* 2]: *L*(*ϕ*(*t*), 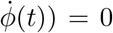. In that case, from Theorem 3the drift of the trajectory is necessarily the drift of a relaxation trajectory between the two timepoints and then, from Corollary 1, ∇_*x*_*V* = 0 on the set of points {*ϕ*(*t*), *t*ℝ^+^}, which is excluded by the condition (C) (when this set is not reduced to a single point). Thus, if *ϕ*(*T*_1_) /= *ϕ*(*T*_2_), we have: *V* (*ϕ*(*T*_1_)) > *V* (*ϕ*(*T*_2_)).

By definition, for any basin *Z*_*i*_ and for all *x* ∈ *Z*_*i*_ there exists a relaxation trajectory connecting *x* to the associated attractor 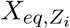. So ∀*x* ∈ *Z*_*i*_, 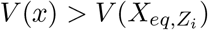.

*Proof of Theorem 4.(iv).* Let *V* be a solution of (25) satisfying the condition (C). We consider trajectories solutions of the system defined by the drift 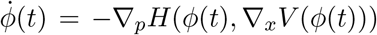. We recall that from (iii), the condition (C) ensures that *V* decreases on these trajectories, and that for all *t*_1_ < *t*_2_: *V* (*ϕ*(*t*_1_)) − *V* (*ϕ*(*t*_2_)) = *Q*(*ϕ*(*t*_2_), *ϕ*(*t*_1_)) > 0. Then, the hypothesis 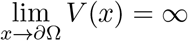 ensures that such trajectories cannot reach the boundary *∂*Ω: if it was the case, we would have a singularity inside Ω, which is excluded by the condition *V* ∈ *C*^1^(Ω, ℝ). The same reasoning also ensures that there is no limit cycle or more complicated orbits for this system.

Recalling that from (i) and (ii), the fixed point of this system are reduced to the points where ∇_*x*_,*V* = 0 on Ω, we conclude that for all *x* ∈ Ω, there exists a fixed point *a* ∈ Ω, satisfying ∇_*x*_*V* (*a*) = 0, such that a trajectory solution of this system converges to *a*, *i.e*: *V* (*x*) − *V* (*a*) = *Q*(*a, x*).

As from the inequality (15), we have for every point *a* the relation *V* (*x*) − *V* (*a*) ≤ *Q*(*a, x*), the previous equality corresponds to a minimum and we obtain the formula:

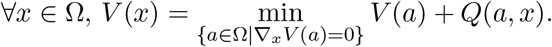

*Proof of Theorem 4.(v).* Let *V* be a solution of (25) satisfying the condition (C). We denote by *ν*_*V*_ the drift of the optimal trajectories *ϕ*_*t*_ on [0, *T*] satisfying the system (16) associated to *V*: ∀*t* ∈ [0, *T*], 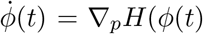, ∇_*x*_*V* (*ϕ*(*t*))) = *ν*_*V*_ (*ϕ*(*t*)). We call trajectories solution of this system *reverse fluctuations* trajectories.

For any basin *Z*_*i*_ associated to the stable equilibrium of the deterministic system 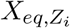, we have:

- From (i), 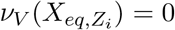 and 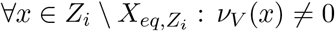.
- From (iii), we know that *V* increases on these trajectories: 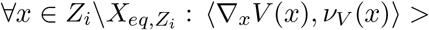 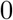.
- From (iii), we also have: 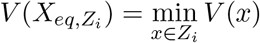.

Without loss of generality (since we only use ∇_*x*_*V*), we can assume 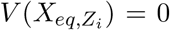. We have then: 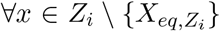, *V* (*x*) > 0. Moreover, since we have assumed that 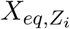 is isolated, there exists *δ*_*V*_ > 0 such that *Z*_*i*_ contains a ball 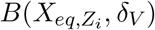. Therefore, *V* reaches a local minimum at 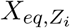. Conversely if *V* reaches a local minimum at a point 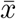, then 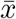 is necessarily an equilibrium (from (ii)), and the fact that *V* strictly decreases on the relaxation trajectories ensures that it is a Lyapunov function for the deterministic system, and then that 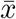 is a stable equilibrium. The stable equilibria of the deterministic system are thereby exactly the local minima of *V*, and for any attractor 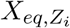, *V* is also a Lyapunov function for the system defined by the drift −*ν*_*V*_, for which 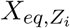 is then a locally asymptotically stable equilibrium. Thereby, stable equilibria of the deterministic system are also stable equilibria of the system defined by the drift −*ν*_*V*_.

It remains to prove that no unstable equilibria the deterministic system is stable for the system defined by the drift −*ν*_*V*_. Let 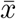 be an unstable equilibrium of the relaxation system, then *V* does not reach a local minimum at 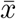. Therefore, as close as we want of 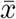 there exists *x* such that 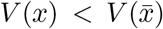. We recall that reverse fluctuations trajectories *ϕ*_*t*_ starting from such a point and remaining in Ω will have *V* (*ϕ*(*t*)) striclty decreasing: by Lyapunov La Salle principle, they shall be attracted towards the set {*y,* ∇〈*V* (*y*), *ν*_*V*_ (*y*)〉 = 0}, which contains from (iii) only the critical points of *V*, which are from (i) the equilibria of both deterministic and reverse fluctuation systems. In particular, either *ϕ*_*t*_ leaves Ω (and the equilibrium is unstable) or *ϕ*_*t*_ converges to another equilibrium (since they are isolated) and this contradicts the stability. So we have proved that stable equilibria of both systems are the same.

We then obtain that for any *Z*_*i*_, there exists *δ*_*V*_ such that:

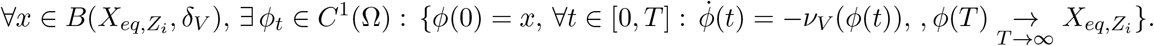

Reverting time, any point of 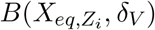 can then be reached from any small neighborhood of 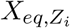. We deduce from Lemma 1 that:

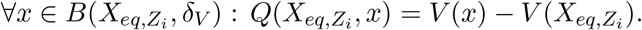

Applying exactly the same reasoning to another function 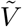 solution of (25) and satisfying (C), this ensures that 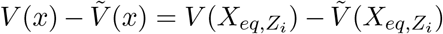, at least for 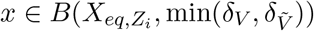. We recall that that from Lemma 2, two optimal trajectories *ϕ*_*t*_, 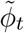 solutions of the system (16), associated respectively to two solutions *V* and 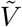 of the equation (25), cannot cross each other without satisfying 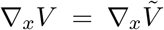 along the whole trajectories. Thereby, we can extend the equality 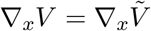 on the basins of attraction associated to the stable equilibrium 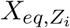 for both systems defined by the drifts −*ν*_*V*_ or 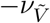. Thus, we have proved that the basins associated to the attractors are the same for both systems. We denote 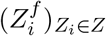 these common basins.

Under the assumption 2. of the theorem, we obtain by continuity of *V* that for every pair of basin 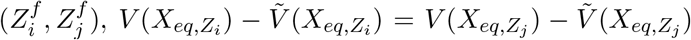. It follows that under this assumption, there exists a constant *c* ∈ ℝ such that for every attractor 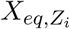:

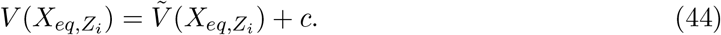

Moreover, the assumption 1. ensures that from Theorem 4.(iv), there exists a fixed point *a*_1_ ∈ Ω (with ∇_*x*_*V* (*a*_1_) = 0), such that a trajectory solution of the system defined by the drift −*ν*_*V*_ converges to *a*_1_, *i.e*: *V* (*x*) − *V* (*a*_1_) = *Q*(*a*_1_*, x*). On one side, if *a*_1_ is unstable, it necessarily exists on any neighborhood of *a*_1_ a point *x*_2_ such that *V* (*x*_2_) < *V* (*a*_1_). As for all *y* ∈ Ω, *Q*(·, *y*) is positive definite, we have then another fixed point *a*_2_ ≠ *a*_1_ such that *V* (*x*_2_) = *Q*(*a*_2_, *x*_2_) + *V* (*a*_2_). We obtain: *V* (*x*) > *h*(*x, a*_1_) + *Q*(*a*_2_, *x*_2_) + *V* (*a*_2_). On the other side, by continuity of the function *Q*(*a*_2_, ·), for every *δ*_1_ > 0, *x*_2_ can be chosen close enough to *a*_1_ such that: *Q*(*a*_2_, *x*_2_) ≥ *Q*(*a*_2_, *a*_1_) − *δ*_1_. We obtain:

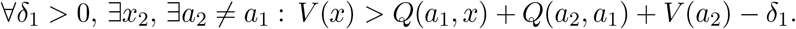

Repeating this procedure until reaching a stable equilibrium at a step *N*, which is necessarily finite because we have by assumption a finite number of fixed points, we obtain the inequality

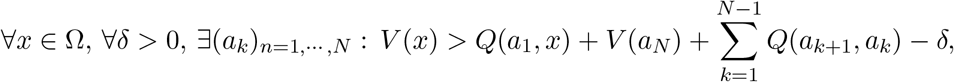

where every *a*_*k*_ denotes a fixed point and *a*_*N*_ is an attractor. Using the triangular inequality satisfied by *Q*, and passing to the limit *δ* → 0, we find that *V* (*x*)−*V* (*a*_*N*_) ≥ *Q*(*a*_*N*_, *x*). Moreover, from the inequality (15), we have necessarily 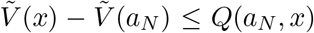. It then follows from (44) that 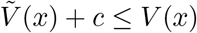.

Applying exactly the same reasoning for building a serie of fixed point 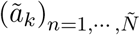 such that 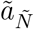 is an attractor and 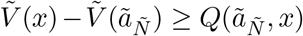, we obtain 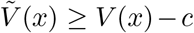. We can conclude:

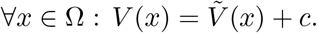

*Proof of Theorem 5.* First, we prove the following lemma:

### Lemma 4.

∀*i* ∈ {1, · · · *, n*}*, we have:*

i. ∃*δ*_*l*_ > 0, ∃*η*_*l*_ > 0, *such that* ∀*x, y* ∈ Ω, if y_i_ < x_i_ ≤ *δ_l_, then we have:*

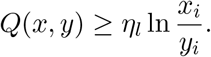
ii. ∃*δ*_*r*_ < 1, ∃*η*_*r*_ > 0*, such that* ∀*x, y* ∈ Ω, if y_i_ > x_i_ ≥ *δ_r_, then we have:*

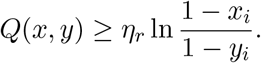

*Proof.* (i) We denote 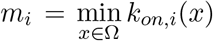. We have *m*_*i*_ > 0 by assumption. We choose a real number *δ* which satisfies these two conditions:

1. 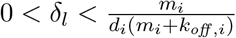,
2. 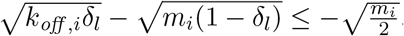.

On the one hand, we recall that the function 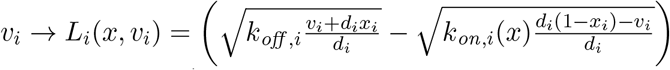 is convex and vanishes only on 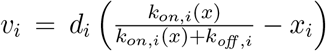. Then, *L*_*i*_(*x,* ·) is decreasing on 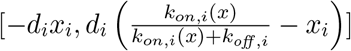.

On the other hand, for all *x* ∈ Ω, if *x*_*i*_ ≤ *δ*_*l*_, we have necessarily: 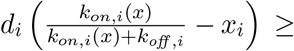 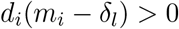 from the condition 1. Then, we obtain that for all *x* ∈ Ω, if *x*_*i*_ ≤ *δ*_*l*_:

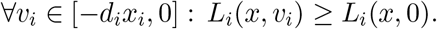

From the condition 2., we also see that for all *x* ∈ Ω, if *x*_*i*_ ≤ *δ*_*l*_:

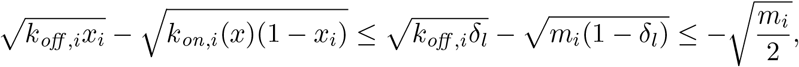

which implies:

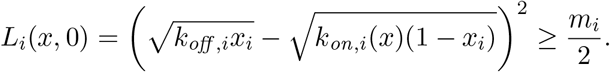

Then, we obtain that for any admissible trajectory 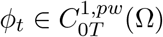 (*i.e* with a velocity in Ω^*v*^(*ϕ*(*t*)) at all time) and such that *ϕ*(0) = *x* and *ϕ*(*T*) = *y*, if *y*_*i*_ < *x*_*i*_ ≤ *δ*_*l*_ we have:

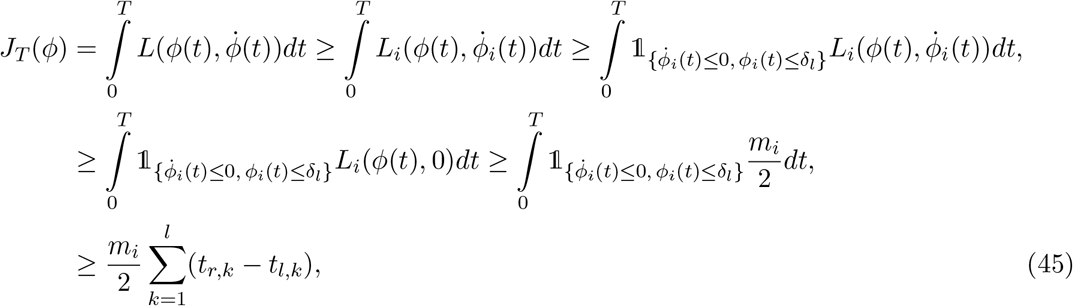

where we denote {[*t*_*l,k*_, *t*_*r,k*_]}_*k*=1, …,l_ the *l* intervals on which the velocity 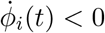 and *ϕ*_*i*_(*t*) < *δ*_*l*_ on the interval [0, *T*]. As we now by assumption that *ϕ*_*i*_(0) = *x*_*i*_ ≤ *δ*_*l*_ and *ϕ*_*i*_(*T*) = *y*_*i*_ < *ϕ*_*i*_(0), this set of intervals cannot be empty.

Moreover, for every *k* = 1, · · ·, *l*, we have by assumption: 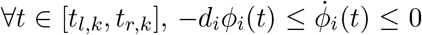.

Then:

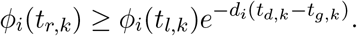

As by definition, for every 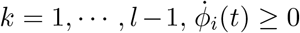 on [*t*_*r,k*_, *t*_*l,k*+1_], we have *ϕ*_*i*_(*t*_*l,k*+1_) ≥ *ϕ*_*i*_(*t*_*r,k*_) (because 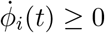 on [*t*_*r,k*_, *t*_*l,k*+1_]). Finally, we obtain:

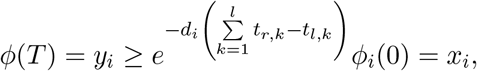

which implies:

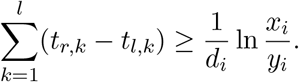

This last inequality combined with (45) allows to conclude:

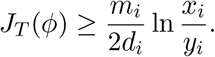

Thus, if *δ*_*l*_ satisfies conditions 1. and 2., and fixing 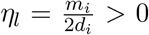, for every *x, y* ∈ Ω such that *y*_*i*_ < *x*_*i*_ ≤ *δ*_*l*_ we have:

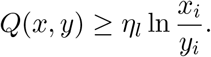

(ii) Denoting 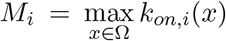 (which exists by assumption), we chose this time the real number *δ*_*r*_ in order to satisfy these these two conditions:

1. 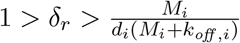,
2. 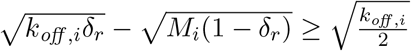,

and we fix 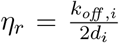. The rest consists in applying exactly the same reasoning than for the proof of (i) in a neighborhood of 1 instead of 0.

We deduce immediately the following corollary:

#### Corollary 2.

∀*x* ∈ Ω, 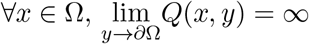.

Let us denote *V* ∈ *C*^1^(Ω, ℝ) a solution of the equation (25), which satisfies the condition (C). From the proof of Theorem 4.(v), we know that for any attractor 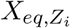, there exists a ball 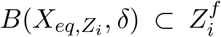, where 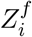 is the basin of attraction of 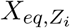 for the system defined by the drift 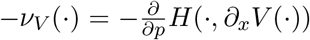. Moreover, as *V* decreases on trajectories solutions of this system, the set 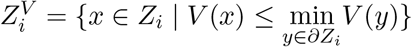 is necessarily stable: we have 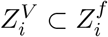.

We deduce that:

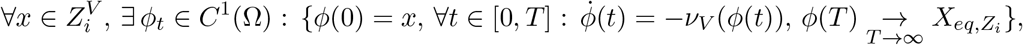

and in that case 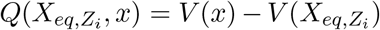. If there existed 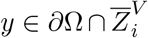, we would have, by continuity of *V* and from Corollary 2:

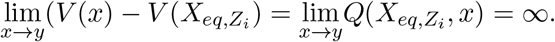

It would imply that 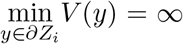, which is impossible when *∂Z*_*i*_ ≠ *∂*Ω, which is necessarily the case when there is more than one attractor.

Thus, there exists at least one point *x*^*i*^ on the boundary *∂Z*_*i*_ \ *∂*Ω, such that for any neighborhood of 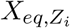, there exists a fluctuation trajectory starting inside and converging to *x*^*i*^.

We recall that we assume that 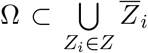. Then we have 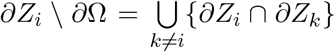 and there exists *Z*_*j*_ such that: *x*^*i*^ ∈ *∂Z*_*i*_ ∩ *∂Z*_*j*_ = *∂Z*_*i*_ \ *R*_*ij*_. We obtain, by continuity of *V*:

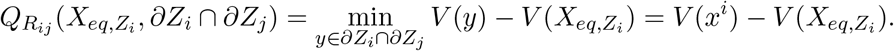

It remains to prove that under the assumption (A) of the theorem, 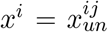. On one hand, from Theorem 4.(iii), *V* decreases on the relaxation trajectories. On the other hand, if every relaxation trajectories starting in *∂Z*_*i*_ ∩ *∂Z*_*j*_ stay inside, they necessarily converge from any point of *∂Z*_*i*_ ∩ *∂Z*_*j*_ to a saddle point (also in *∂Z*_*i*_ ∩ *∂Z*_*j*_). Then, the minimum of *V* in *∂Z*_*i*_ ∩ *∂Z*_*j*_ is reached on the minimum of *V* on 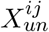 (the set of all the saddle points in *∂Z*_*i*_ ∩ *∂Z*_*j*_), which is 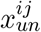. Thus, if every relaxation trajectories starting in *∂Z*_*i*_ ∩ *∂Z*_*j*_ stay inside, then *X*_*Z*_ ∈ *∂Z*_*j*_ implies 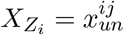. The theorem is proved.

## I Algorithm to find the saddle points

We develop a simple algorithm using the Lagrangian associated to the fluctuation trajectories (28) to find the saddle points of the deterministic system (4). This Lagrangian is a nonnegative function which vanishes only at the equilibria of this system. Then, if there exists a saddle point connecting two attractors, this function will vanish at this point. Starting on a small neighborhood of the first attractor, we follow the direction of the second one until reaching a maximum on the line (see 15a). Then, we follow different lines, in the direction of each other attractor for which the Lagrangian function decreases (at least, the direction of the second attractor (see 15b)), until reaching a local minimum. We then apply a gradient descent to find a local minimum (see 15c). If this minimum is equal to 0, this is a saddle point, if not we repeat the algorithm from this local minimum until reaching a saddle point or an attractor. Repeating this operation for any ordered couple of attractors 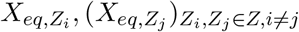, we are likely to find most of the saddle points of the system. This method is described in pseudo-code in Algorithm 1.

**Figure 15:**
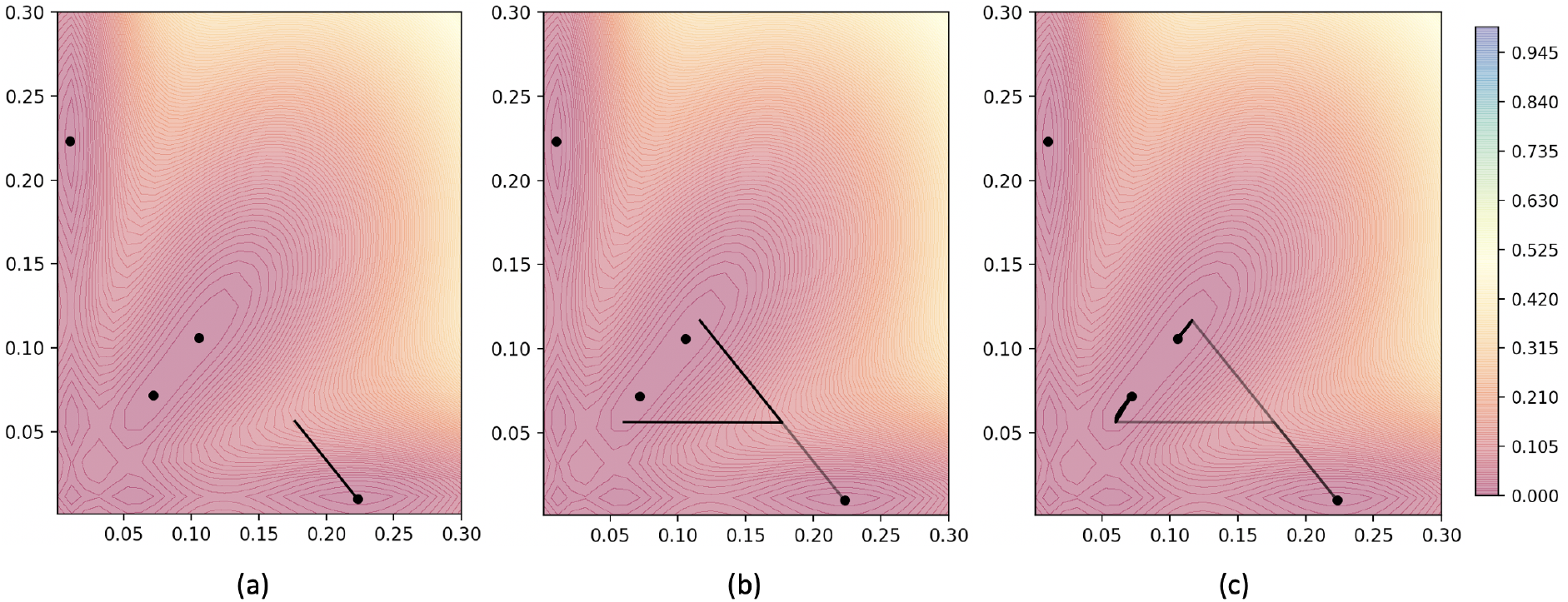
Saddle-point algorithm between two attractors. The color map corresponds to the Lagrangian function associated to the fluctuation trajectories.

### Algorithm 1

Find the list of saddle points: list-saddle-points

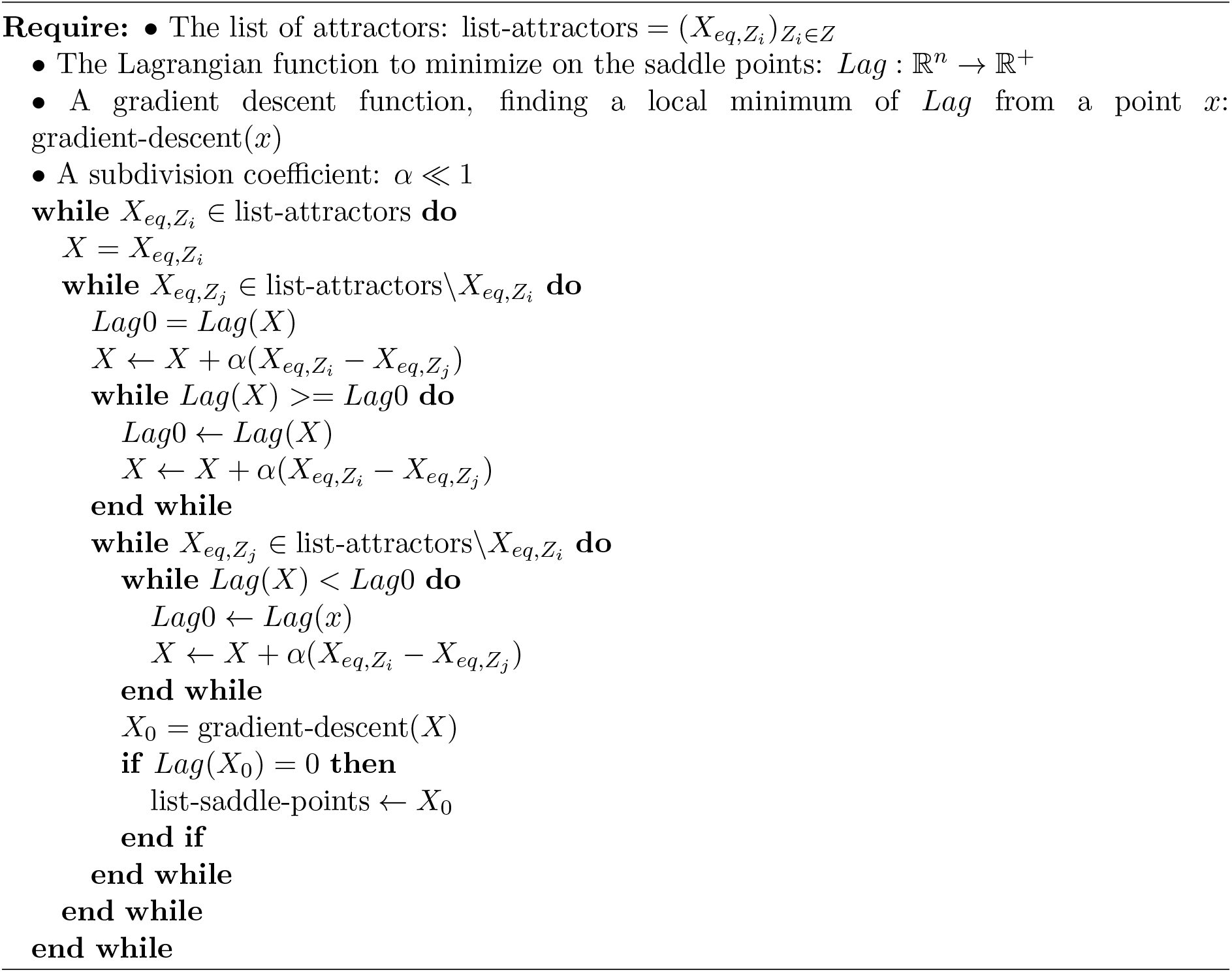

## J Applicability of the method for non-symmetric networks and more than two genes

We present in Figures 16b and 17b an analogy of Figure 8, which was presented for the toggle-switch network, for two non-symmetric networks of respectively 3 and 4 genes. The networks are presented on the left-hand side of Figures 16b and 17b: the red arrows represent the inhibitions and the green arrows represent the activations between genes. A typical random trajectory for each network is presented in Figures 16a and 17a.

We recall that we build the LDP approximation (in red) by using the cost of the trajectories satisfying the system (28) between the attractors and the saddle points of the system (4). The cost of these trajectories is known to be optimal when there exists a solution *V* of the equation (25) which verifies the relations (26), which can generally happen only under symmetry conditions. This is not the case nor for the 3 genes network of Figure 16b when there is no symmetry between the interactions, neither for the 4 genes network of Figure 17b. Then, we could expect that these LDP approximations would be far from the Monte-Carlo and AMS computations, especially for the 4 genes network, since we have no symmetry between the interactions, not only in value but also in sign. However, we observe that the approximations given by our method seem to remain relatively accurate.

**Figure 16:**
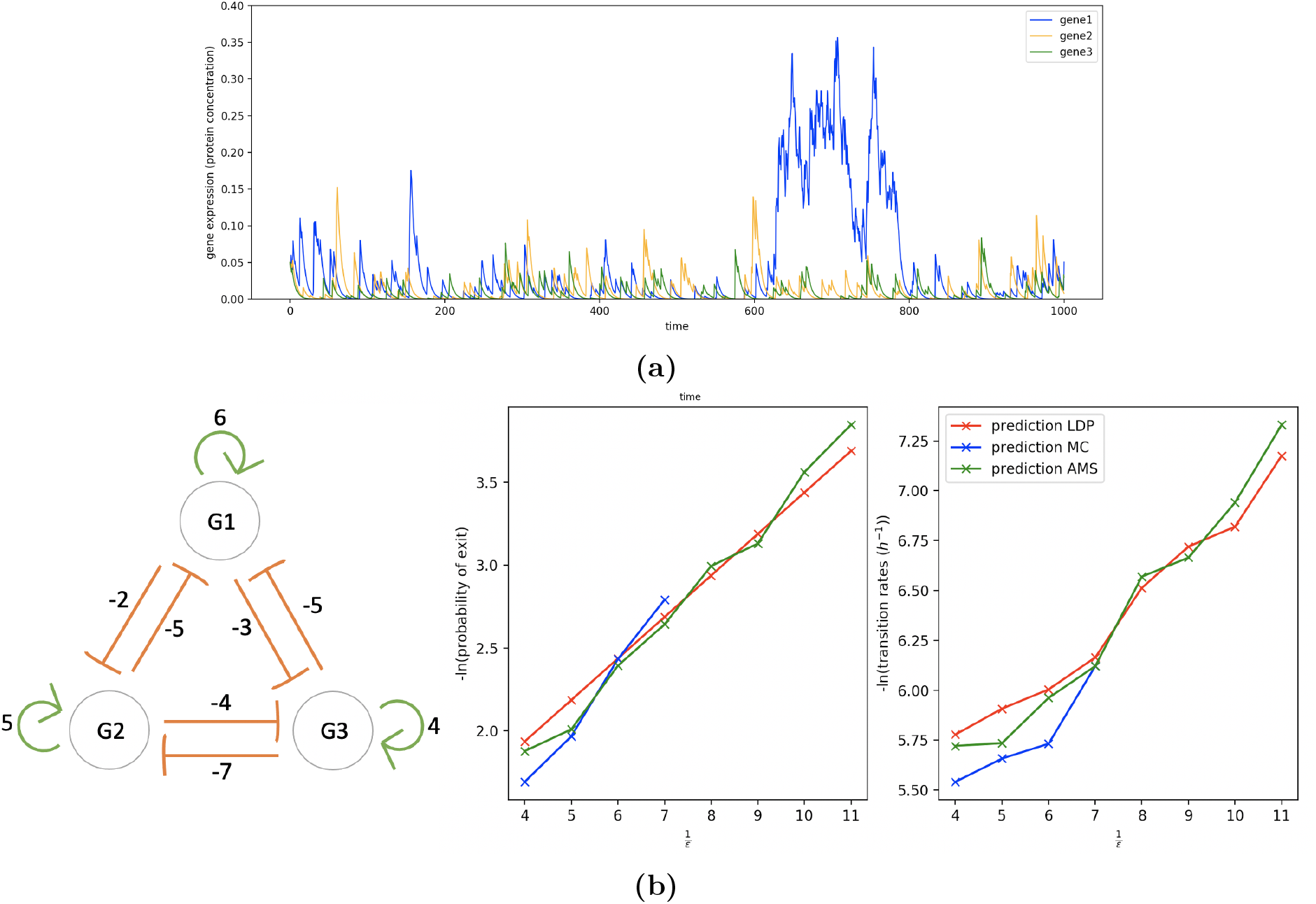
16a: A random trajectory associated to the non-symmetric toggle-switch network of 3 genes with *ε* = 1/8. The network is associated to 2 attractors only: *Z*_−−−_ all the genes inactive, and *Z*_+−−_ gene 1 active and gene 2 and gene 3 inactive due to the inhibitions. 16b: Analogy of Figure 8 between *Z*_−−−_ and *Z*_+−−_. We see that the analytical approximations of the transition rates are very accurate although we had no theoretical evidence for the trajectories computed by the method presented in Section 4.3 to be optimal.

**Figure 17:**
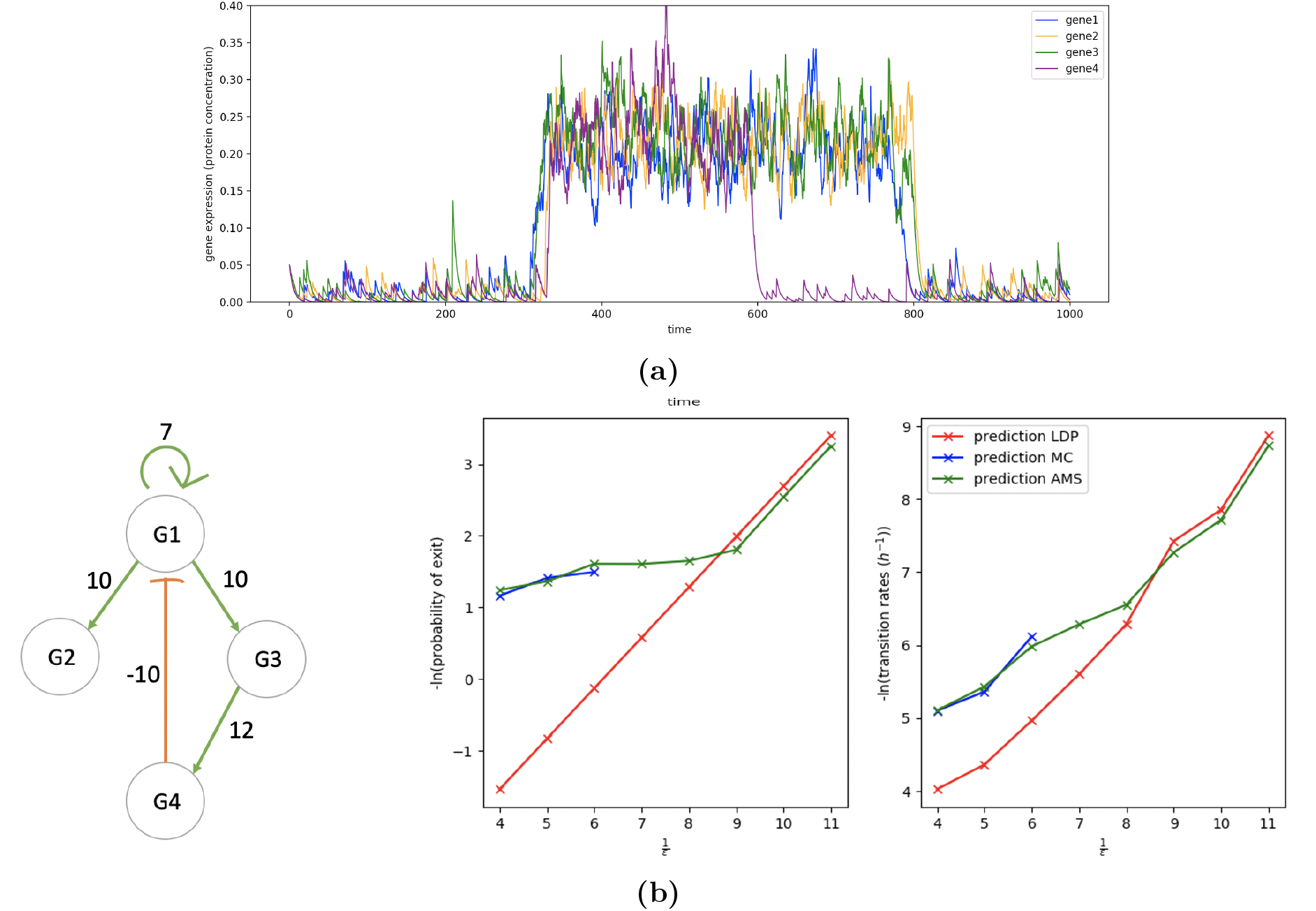
17a: A random trajectory associated to the non-symmetric 4 genes network with *ε* = 1/8. The network is associated to 3 attractors: *Z*_−−−−_ all the genes inactive, *Z*_+++−_ genes 1-2-3 active and gene 4 inactive, and *Z*_++++_ all the genes active. 17b: Analogy of Figure 8 between *Z*_++++_ and *Z*_+++−_. The analytical approximations seem to become accurate from *ε* ≃ 1/9.

